# Three nitrogen atoms turn old kinase inhibitors into new targetable remedy

**DOI:** 10.64898/2026.04.11.717649

**Authors:** Balázs Besztercei, Réka Antal, Leena Tähtivaara, Birgitta Lappeteläinen, Niina Jääskeläinen, Attila Szappanos, Ákos Lukáts, Alexandra Pál-Kajtár, András Budai, Marc Cerrada-Gimenez, Krisztián A. Kovács

**Affiliations:** Semmelweis University, Institute of Translational Medicine; Translational Retina Research Group. Tűzoltó u. 37-47., 1094 Budapest, Hungary; Experimentica Ltd., 1 Microkatu, 70210 Kuopio, Finland; Semmelweis University, Department of Pathology, Forensic and Insurance Medicine. Üllői út. 93., 1091 Budapest, Hungary

**Keywords:** Neovascularization, age-related macular degeneration, diabetic retinopathy, tyrosine kinases, drug targeting, photoactivable inhibitor

## Abstract

Pathological neovascularization in the eye is a significant contributor to vision loss in diseases such as age-related macular degeneration (AMD) and diabetic retinopathy. While anti-VEGF biologics are effective, they require repeated intravitreal injections and carry procedural risks. Here, we report a novel principle for designing photoactivatable VEGFR2 inhibitors, along with two examples, EYE1090 and EYE1118, engineered from the sunitinib and vorolanib scaffolds, respectively. Azido-functionalization in these molecules enables light-triggered receptor binding while preserving potent inhibition in the dark. Both compounds exhibit significantly enhanced activity upon exposure to green light capable of reaching the retina, also in the elderly. *In vitro,* the compounds robustly inhibited angiogenesis and endothelial migration that was further potentiated by light. *In vivo,* orally administered EYE1090 and EYE1118 suppressed VEGF-induced retinal leakage and reduced lesion size in a mouse model of choroidal neovascularization at doses tenfold lower than parental compounds. Photoactivation also influenced compound biodistribution, suggesting light-guided targeting. Acute toxicity studies revealed no hepatotoxicity. This strategy exploits the natural light-focusing anatomy of the eye to locally activate systemically administered drugs, thereby reducing therapeutic doses and systemic exposure. Our findings introduce a light-targeted pharmacological approach for treating retinal diseases using photoactivable kinase inhibitors.

## INTRODUCTION

Neovascularization (*1*) can be traced back to the pathological activation of the VEGF-VEGFR2 axis and leads to the formation of a superfluous amount of ectopic neovessels. Since solid tumors require oxygen and nutrients to grow, this process has been considered the Achilles’ heel of tumorigenesis. In line with this observation, sunitinib, a potent angiokinase inhibitor, has been approved for the treatment of three different types of cancer (*2*). The molecular mechanisms underlying choroidal and retinal neovascularization are highly similar to those in tumors; however, treatments for ocular diseases must meet stricter safety standards. Correspondingly, vorolanib, despite its superior safety profile to sunitinib and more favorable kinase specificity, has failed in human clinical trials in orally administered patients with age-related macular degeneration due to its dangerous hepatobiliary effects (*3,4*).

At present, pharmacological suppression of ocular neovascularization relies exclusively on intravitreal injections, which are associated with serious adverse events and represent a major burden for the patients. The gold standard treatments for age-related macular degeneration (*5,6*) and diabetic retinopathy (*7*) are based on anti-VEGF biologics such as aflibercept (Eylea). Moreover, several clinical trials have recently been conducted to introduce biosimilars of aflibercept onto the market, and further macromolecules working along the same principles (*8*). Some progress at the conceptual level is indicated by the fact that implanted ocular depot formulations of vorolanib are being tested and show promising efficacy in human clinical trials. Nonetheless, the best outcome the patients can hope for from these investigations is a reduced frequency of injections, given that the implant is also delivered intravitreally.

A largely overlooked strategy involves small-molecule receptor tyrosine kinase inhibitors, which can effectively suppress neovascularization and, in principle, be administered orally. However, their use remains limited to oncology (*9*) due to frequent and serious systemic side effects.

In the present work, we uncover a novel, light-mediated strategy to target orally taken oxindole-based angiokinase inhibitors directly to the tissue to be treated so that the therapeutic effect can be exerted locally. According to our expectation, this approach would allow for a drastic reduction of the serum concentrations required for treating ocular diseases and may also serve as a light-targeted therapy in oncology. We show that substantially lower doses of our new compounds can effectively reduce choroidal neovascularization in mice and VEGF-induced leakage of retinal capillaries in rats. Preliminary hepatotoxicity data also suggest a favorable safety profile. Beyond ophthalmology, this targeted method may offer a new modality for treating cancer with reduced systemic toxicity.

## MATERIALS AND METHODS

### Cellular viability assay

On day 1, 30,000 NIH/3T3 cells were seeded in each well of a 96-well plate in 180-180 μl of DMEM (Gibco, Cat. No.: 11995065) containing 10% of FBS and 1% of penicillin/streptomycin, and incubated for 16 hours (37, 5% CO_2_). On day 2, cells were treated with EYE1118, EYE1090, vorolanib (MedChemExpress LLC), or sunitinib (MedChemExpress LLC) at the indicated concentrations (0.1 – 100 μM) while the plates were protected from light. In the case of controls, cells were treated with 0.2% DMSO, 2% DMSO, 50% Geneticin, or media alone. The plates were incubated for 24 hours at 37 and 5% CO_2_ in the dark. On day 3, alamarBlue (Invitrogen, Cat. No.: DAL1025) was added to each well while plates were kept in the dark. After 5 hours of incubation, the relative fluorescence was measured using a CLARIOstar (BMG Labtech) plate reader set at an excitation wavelength of 570 nm and emission wavelength of 610 nm. The mean and SD values of viability percentage were calculated using the GraphPad Prism software. Data were analyzed by two-way ANOVA.

### Luciferase measurements

#### Cells

Commercially available (BPS BioScience Inc., Cat. No.: 79387) VEGFR2 / NFAT Reporter - HEK293 recombinant cell line was used to measure the inhibition by EYE1090 and EYE1118 according to the manufacturer’s instructions.

#### Seeding

On the day preceding the measurement, 50,000 recombinant HEK cells were seeded into each well of a 96-well plate in a way that the experiments could be conducted in triplicate. On the day of the measurement, the growth medium (BPS BioScience Inc., Cat. No.: 79528) was replaced with assay medium (BPS BioScience Inc., Cat. No.: 60187-1), and then the confluence was verified. Only plates with 100% confluence in each well were processed further.For data presented in one graph (one panel of Figure 2) only cells from the same seeding procedure were used to ensure the uniform expression of the NFAT-Luc and CAG-VEGFR2 constructs.

#### Measurement

Inhibitor molecules were applied in the indicated concentration and were left on the cells for 50 minutes to allow for the inward diffusion. For the conditions requiring irradiation, the plate was then placed under an LED light source within the incubator (37°C, 5% CO_2_) for the last 10 minutes of the 1-hour-long incubation with the inhibitors, while the other plate was kept in the dark (also at 37 °C, 5% CO_2_). Following the pre-incubation with inhibitors, we treated the cells for 4 hours with VEGF without removing the inhibitors. For this purpose, we used 50 ng/ml of commercially available human VEGF-165 (Sf9 derived, from BPS BioScience Inc., Cat. No.: 91001-1) added directly into the cell-culture medium. After completion of all the incubation steps, the luciferase activity was measured using the commercially available “ONE-step luciferase assay system” (BPS BioScience Inc., Cat. No.: 60690-2) and a CLARIOstar (BMG Labtech) plate reader that quantified luminescence originating from each well of the 96-well plate. The IC_50_ values were calculated using the GraphPad Prism software.

### In vitro angiogenesis

To test the inhibitory effect of the compounds with light and without light on in vitro angiogenesis (hereinafter referred to as “tubulogenesis”), we have used human retinal microvascular endothelial cells (HRMEC) from Cell Systems (Kirkland, WA 98033, cat. N°: ACBRI 181). VEGF-165 (Invitrogen) was only used as a positive control in these experiments, otherwise tubulogenesis relied on endogenous VEGF produced by the culture.

Ninety-six well plates were coated with Geltrex basement membrane matrix with reduced growth factor content (Gibco), centrifuged at 4 °C, and incubated in a humidified incubator at 37°C for 30 minutes. Before seeding, HRMEC cells were treated with compounds (0.1 nM - 10 µM) in EBM-2 basal medium (Lonza) in the dark for 10 minutes. Next, cells were seeded onto the Geltrex matrix (1×103 cells per well) and then exposed to light (cold white LED, 693 lux) or kept in the dark (control plate) for 10 minutes at 37 °C and 5% CO2. After 12 hours of incubation at 37 °C and 5% CO2, cells were stained with Calcein AM(Invitrogen) for 30 minutes in the dark. A series of images spanning 120 µm range along the z axis were acquired and the built-in algorithms called Extended Depth of Field (EDF) create one two-dimensional focused image. All EDF images were analyzed using the freely customizable NIS-Elements General Analysis 3 module. We set up the experiment routine to segment and identify tubular and nodular structures and extracted tube number, node number and the number of meshes as readouts. IC_50_ values were calculated using GraphPad Prism software.

### Endothelial migration assay

On day 1, HRMEC from Cell Systems (Kirkland, WA 98033, cat. N°: ACBRI 181) were serum starved overnight by removing the EGM-2 MV (complete growth medium, Lonza), adding EBM-2 (basal medium, Lonza) containing 0.5% FBS to the cells, and incubating for 12 hours. At the same time, the transwell inserts were coated with collagen I solution (10 µg/cm2) and the plate was incubated overnight at 4 °C.

The next day, the excess collagen coating solution was removed from the inserts, which were then washed with 1xPBS, and their basolateral surface was coated with 0,5% gelatin solution for 30 min at room temperature. Finally, the inserts were washed with EBM-2 medium.

At the same time, the inhibitors were diluted at different concentrations (0,001-10 µM) with EBM-2 medium. Cells were then harvested using Accutase solution (Merck), their number was determined, and a suspension with an appropriate cell number was prepared with EBM-2 medium. Next, the inhibitors were mixed with the cells and incubated for 10 minutes in the incubator. Then one plate was exposed to light (cold white LED, 693 lux) or kept in the dark (control plate) for 10 minutes at 37 °C and 5% CO2. The cells were then transferred to the transwell insert. Fifty ng/ml VEGF or vehicle control (EBM-2 basal medium) was added to the lower chamber. Plates were incubated for 6 hours at 37 C and 5% CO2 in the dark. Next, the medium was aspirated from the lower and upper chambers, and the cells were washed with 1x PBS and fixed with 4% PFA for 10 min. The plate was washed with 1x PBS and nuclei were stained with 10µg/ml Hoechst 33342 dye (Invitrogen) for 10 min. Before imaging the non-migrated cells were scraped with pipette tips and the remaining cells on the basolateral surface (migrated cells) were imaged using an epifluorescence microscope (Nikon). Images were captured using a DAPI filter set and 4x/0.13 and 10x/0.4 objectives. For each well, 2×2 large images were taken in order to capture the entire surface of the membrane. Cell nuclei were counted using Nis-Elements software. IC_50_ values were calculated using GraphPad Prism software.

### Pharmacokinetic studies of EYE1090 photoactivation in BALB/c mice

#### Animals

Male Balb/cJRj mice 8 weeks old at the start of the study (Janvier, France) were housed in individually ventilated cages (IVC) with aspen bedding, nesting material and gnawing blocks (Populus tremula, Tapvei Estonia OÜ) and polycarbonate igloos as enrichment, at a constant temperature (22 ± 1 °C), relative humidity (50 ± 10 %) and in a light controlled environment (lights on from 7 am to 7 pm) (300 lux). Mice had ad libitum access to food (Rat/Mouse maintenance V1534-000, ssniff Spezialdiäten GmbH) and tap water. Experiments started after a three-week quarantine and acclimatization in the vivarium. 48 h prior to the treatment administrations, all the mice were moved into a dim red light-controlled environment. All animals were treated in accordance with the ARVO Statement for the Use of Animals in Ophthalmic and Vision Research and the EC Directive 2010/63/EU for animal experiments, using protocols approved and monitored by the Animal Experiment Board of Finland (Experimentica Ltd. animal license number ESAVI-11107-2023).

#### Compound administration

In order to enhance the transport of the new chemical entities through the blood-retina-barrier, the third-generation P-gp inhibitor elacridar was administered intragastrically (i.g.) to all the mice 8 h prior to the treatment. The new chemical entities were administered at 40 mg/kg i.g. Following this procedure, all the animals were kept under dim light conditions (red light 5 lux) for 30 min. Afterwards, the bright light groups were placed under a 1,000 lux white light environment for an additional 3.5 h, while the rest of the animals were kept under dim light conditions. Four hours after administration, the mice were sacrificed under red light conditions. Plasma and cellular debris for PK analysis were collected under red light conditions.

#### Sample preparation for PK analysis

The extraction procedure was performed under minimal exposure to light. The plasma samples were stored at −80, and before the extraction procedure, the samples were thawed at room temperature (RT). Plasma samples were placed into 15 mL centrifuge tubes, then 200 µL of 10% NH4OH solution was added to them and vortexed for 30 s. After that, the samples were adjusted with ethyl acetate to 5 mL and vortexed for 30 s and were subsequently centrifuged at 3500 rpm for 10 min at RT. 4 mL of supernatant was removed and placed in a round-bottom flask. The remaining pellet was adjusted with ethyl acetate to 5 mL, then vortexed for 30 s and centrifuged at 3500 rpm for 10 min. 4 mL of the supernatant was removed and placed into the same round-bottom flask with the supernatant from the first round of extraction. The samples were evaporated to dryness by a rotary evaporator. The dried samples were reconstituted with 300 µL of 90% acetonitrile (ACN) in ultrapure water, then dissolved, vortexed, and centrifuged at 10,000 rpm for 10 min at RT. Finally, 200 µL of supernatant was placed into a new amber glass vial.

#### HPLC instrument and measurement conditions

For the HPLC analysis of the samples, the Waters™ e2695 Separation Module (Waters, MA, USA) was used with the Waters™ 2996 PhotoDiode Array Detector (Waters, MA, USA). The chromatographic separation was carried out with the Kinetex 2.6 µm HILIC 100 Å column (Phenomenex, CA, USA). A mixture of „A”: 10% 50 mM ammonium formate and 40% dH2O in ACN, „B”: 10% 50 mM ammonium formate in ACN was used as the mobile phase. The temperature of the chromatographic column was 25 during the runtime. The gradient program is shown in the table below. The UV spectrum of the samples was determined with a diode array detector scan mode (190-600 nm), and EYE1090 was monitored at 444 nm.

**Table 1.** The composition of the mobile phase during gradient elution to recover EYE1090 from the Kinetex column. The indicated amount of the indicated components (**A, B**) were mixed and fed onto the column. **A**: 10% 50 mM ammonium formate and 40% dH_2_O in ACN; **B:** 10% 50 mM ammonium formate in ACN.

| Time (min) | Flow rate (ml/min) | Percentage of A | Percentage of B |
| --- | --- | --- | --- |
| 0 | 0.4 | 10 | 90 |
| 2.5 | 0.4 | 10 | 90 |
| 8 | 0.4 | 50 | 50 |
| 10.5 | 0.4 | 50 | 50 |
| 11.5 | 0.4 | 10 | 90 |
| 14 | 0.4 | 10 | 90 |

### Hepatotoxicity assessment in BALB/c mice

#### Animals

Male and female Balb/cJRj mice 6-10 weeks old at the start of the study (Janvier, France) are housed in individually ventilated cages (IVC) with aspen bedding, nesting material and gnawing blocks (Populus tremula, Tapvei Estonia OÜ) and transparent polycarbonate 130 mm tubes (Datesand group) as enrichment, at a constant temperature (22 ± 1 °C), relative humidity (50±10 %) and in a light controlled environment (lights on from 7 am to 7 pm), either under normal (300 lux) or red light (5 lux) light conditions. Mice will have ad libitum access to food (Rat/Mouse maintenance V1534-000, ssniff Spezialdiäten GmbH) and tap water. Experiments start after a minimum of one week of quarantine and acclimatization in the vivarium. All animals are treated in accordance with the ARVO Statement for the Use of Animals in Ophthalmic and Vision Research and the EC Directive 2010/63/EU for animal experiments, using protocols approved and monitored by the Animal Experiment Board of Finland (Experimentica Ltd. animal license number ESAVI-10546-2023)

#### Compound administration

In order to enhance the transport of the new chemical entities through the blood-retina-barrier, the third-generation P-gp inhibitor elacridar was administered intragastrically (i.g.) to all the mice 8 h prior to the treatment. Both EYE1118 and EYE1090 were administered at 40 mg/kg i.g. via oral gavage; naive animals received the equivalent amount of vehicle per os.

#### Sample collection

Seventy-two hours after compound or vehicle administration, the animals were sacrificed by cardiac puncture, under the surgical plane of anaesthesia. The liver left lobules were collected, fixed in formalin overnight at +4°C. The following day, the tissue samples were dehydrated by placing them in solutions of increasing alcohol concentration. Finally, the tissues were incubated in Xylene prior to being embedded in paraffin. The tissue blocks were stored at RT until analysis.

#### Histology

FFPE samples were cut to 4 μm thick histological slides, then were deparaffinized with Xylol and rehydrated in a graded alcohol series. Standard hematoxylin and eosin staining was performed for general histological evaluation. In addition to the customary application of hematoxylin-eosin staining on the specimens for visualization, picrosirius red staining was employed to unveil fibrotic changes. Finally, PAS and diastase digested PAS reactions were used to evaluate glycogen and glycoprotein accumulation.

### Assessing the retinal permeability in rats

All procedures were performed in concordance with the Association for Research in Vision and Ophthalmology statement for the Use of Animals in Ophthalmic and Vision Research (ARVO statement) and were approved by the Ethics Committee of Semmelweis University and by the Animal Health and Animal Welfare Directorate of the National Food Chain Safety Office (PE/EA/01261-6/2022).

Male Long Evans rats (200-250 g, from Janvier Labs) were used in the study. Animals were housed in a room with constant temperature (22 ± 2 °C) under a 12-12 h alternating light-dark cycle. Standard chow and water were supplied ad libitum. Animals were randomly enrolled into study and control groups after a 1-week accommodation period.

Vascular Endothelial Growth Factor (VEGF, Human VEGF 165 Recombinant Protein, Thermo Fisher) at 50 ng/eye, or sterile saline solution (0.9% NaCl) was administered through the same procedure. The pupils were dilated with Tropicamide eyedrops (5 mg/ml, S.C. Rompharm Company S.R.L.) prior to all procedures and after local disinfection (Povidone Iodine, Betadine eye drops), injections were carried out in general anesthesia (3% isoflurane in 100% oxygen), with a Hamilton glass syringe and a 34-gauge needle (CP-Analitika Kft.) in a final volume of 5 µl/eye. To reduce the risk of post-injection infection, antibiotic eyedrops were also routinely applied (ofloxacin, Floxal 3 mg/ml, Dr. Gerhard Mann - Chemisch-Pharmazeutische Fabrik). In the rare case of intraocular infections, bleeding or retinal detachment, the eyes were excluded from the study.

To examine the effect of EYE 1090 and EYE 1118 on VEGF-induced vascular leakage, animals were randomly divided into 6 groups (n≥7/group). The left eyes received VEGF while the right eyes received vehicle only in all cases. The animals were either treated intragastrically with EYE 1090 (1 mg/kg) or EYE 1118 (1mg/kg) or received vehicle (isotonic solution with equivalent amount of DMSO) for three consecutive days. VEGF or vehicle were injected intravitreally on the third day of treatment.

Vascular leakage measurements were performed 24 hours after the VEGF injections in general anesthesia (1-2% isoflurane in 100% oxygen) following a slightly modified Evans Blue (EB) technique. A 30 mg/ml EB (Sigma-Aldrich Kft.) solution was prepared by dissolving the dye in sterile saline. After sonification and filtration, a dose of 45 mg/kg was injected to each animal through the femoral vein. EB concentrations were estimated from cardiac puncture immediately before perfusion. Blood samples were centrifuged at 12000 rpm for 15 minutes, then the plasma was diluted 1:10000 in deionized formamide (Life Technologies Kft.). Two hours after the EB injection the animals were perfused intracardially with warm heparinized saline solution. The eyes were immediately removed, the retinae were detached, thoroughly dried and weighted. The dye was extracted by incubating in 150 µl formamide for 18 hours at 70 °C.

The EB concentrations of the samples were determined by the fluorescent method. The samples were centrifuged (14000 rpm, 24 °C, 60 minutes) and 100 µl supernatant was measured on a CLARIOstar plate reader (BMG Labtech) The excitation wavelength was 620 nm, and the measurement was performed at 680 nm. A standard dilution series was used to establish a calibration curve and describe the concentration emission relationship. Linear regression was used to create an EB calibration curve and fit an equation that was used to determine retinal EB concentrations. As retinas also have autofluorescence at the wavelength used for measurements, all EB values were corrected for the retinal background fluorescence determined in an independent experiment.

Vascular leakage was calculated using the following equation:

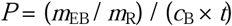

Where *P* stands for permeability, *m*_EB_ is the mass of EB in the retina (determined using the calibration curve), *m*_R_ is the dry retinal weight, *c*_B_ is the EB concentration in the plasma, and *t* is the time during which EB was present in the circulation. Permeability values were expressed in µl × g^−1^ × h^−1^ units.

### Assessing the efficacy of EYE1090 and EYE1118 in a mouse CNV model

Adult C57BL/6JRj mice (9 weeks old, Janvier Labs) were maintained on a 12-h light/dark cycle and provided standard food (Rat/Mouse maintenance V1534-703, irradiated, sniff Spezialdiäten GmbH) and water ad libitum. All experiments were performed in accordance with the ARVO Statement for the Use of Animals in Ophthalmic and Vision Research and the EC Directive 2010/63/EU for animal experiments, using protocols approved and monitored by the Project Authorisation Board of Finland.

EYE1118 and EYE1090 were administered at a dose of 1 mg/kg, or vehicle (10% DMSO) was administered intragastrically to the mice once daily for three days, starting two days before CNV induction. All treatments were administered under red light, and animals were kept in the dark for an additional 30 minutes before being transferred into normal light conditions.

To induce choroidal neovascularization (CNV), the mice were anesthetized with a mixture of ketamine and medetomidine hydrochloride (30 and 0.4 mg/kg, respectively), and the pupils were dilated with a drop of 0.5% tropicamide. Three laser spots surrounding the optic nerve were created using a green diode laser (spot size 100 µm, duration 120 ms, power 130 mW, Oculight TX. Iridex Corp.). The formation of a bubble during lasering and perforation of the Bruch’s membrane, seen in spectral-domain optical coherence tomography (SD-OCT), indicated a successful induction.

Choroidal vascular leakage was assessed by indocyanine green angiography (ICGA) using a Heidelberg Spectralis HRA2 system (Heidelberg Engineering). The mice were administered 130 µl of indocyanine green (2 mg/ml) subcutaneously, with images of the choroid and retina acquired immediately and at 60-second intervals for 5 minutes. ICGA was performed immediately after lasering (day 0), and on days 3, 5, and 7. ICGA positive area was manually thresholded using the FIJI image analysis suite (ImageJ v.1.53t). The areas were normalized to the Day 3 results.

### Immunoblotting

Serum-starved VEGFR2/NFAT reporter HEK293 cells (BPS BioScience Inc., Cat. No.: 79387; 500.000 cells/well) were incubated in 6-well plates with EYE1090 and EYE1118 (100, 10, 50, 1 nM) in the dark. After 1 hour of incubation, one plate was irradiated with 693 lux cold white LED light source for 10 minutes, and the control plate was kept in the dark. Cells were stimulated with 10 ng/mL of VEGF-165(Invitrogen) or vehicle control for the desired period. The plates were then placed on ice, and the cells were washed twice with ice-cold PBS and lysed in RIPA lysis buffer containing complete protease inhibitor, 1mM PMSF, and 1mM Na3VO4. The lysates were incubated at 4 C for 30 min and centrifuged at 12000g for 5 min at 4 C. The supernatant was transferred to a clean 1.5 ml tube, and the protein concentration was determined using Pierce™ Rapid Gold BCA Protein Assay Kit(Thermo Scientific). Equal amounts of protein (20µg) from cell lysates were analyzed by SDS-PAGE followed by Western blot to detect the levels of phosphorylated VEGFR2, total VEGFR2, and β-actin. The following primary antibodies were used: pY1175-VEGFR2 (Thermo Fisher), pY951-VEGFR2 (Thermo Fisher), VEGFR2 (B.309.4, Thermo Fisher), and β-actin (5J11, Invitrogen). PVDF membranes were incubated with primary antibodies at 4 C overnight and followed by secondary antibodies, HRP-conjugated goat anti-Rabbit and anti-Mouse (Invitrogen) for 1 hour at RT. Chemiluminescent signals were captured using a CCD camera-based imager (VWR International). After detecting the level of phosphorylated receptors, the membranes were stripped and reprobed with primary antibody against the total level of VEGFR2 (B.309.4). Beta-actin was used as a loading control.

### Synthesis of EYE1090

EYE1090 stands for 5-[(Z)-(5-azido-6-chloro-2-oxo-1,2-dihydro-3H-indol-3-ylidene) methyl]-N-[2-(diethylamino)ethyl]-2,4-dimethyl-1H-pyrrole-3-carboxamide (C_22_H_26_ClN_7_O_2_) that was synthetized according to the procedure described in detail in our previous patent (*10*). Purity was as indicated in the patent (*10*)

### Synthesis of EYE1118

EYE1118 stands for 5-[(Z)-(5-azido-6-chloro-2-oxo-1,2-dihydro-3H-indol-3-ylidene) methyl]-N-[(3S)-1-(dimethylcarbamoyl) pyrrolidin-3-yl]-2,4-dimethyl-1H-pyrrole-3-carboxamide (C_23_H_25_ClN_8_O_3_) that was synthetized according to the procedure described in detail in our previous patent (*10*). Purity was as indicated in the patent (*10*)

### Statistical analysis

For permeability tests in rats, data were expressed as mean±SD. For comparing permeability values, normality was checked with Shapiro-Wilk’s test, and the values were compared using one-way ANOVA and Tukey’s post-hoc test. The significance level was set at P<0.05.

To analyze the severity of the CNV in the mouse, data were analyzed by Mixed-effects analysis, and differences from the Vehicle group were analyzed by Holm-Šidák’s multiple comparison test. Data were graphed and statistically analyzed using the GraphPad Prism software (v.10.4.0. GraphPad Software LLC).

## RESULTS

### Azido-functionalization of oxidole-based VEGFR2 inhibitors

In a previous patent, we have shown that existing VEGFR2 inhibitors can be turned into photoactivable molecules via azidation and these new chemical entities show increased potency upon illumination (*11*), presumably via a covalent bond to the receptor. This finding is particularly relevant for ophthalmic indications, such as age-related macular degeneration (AMD), diabetic retinopathy, and diabetic macular edema, where the activating light can naturally reach the retina without invasive intervention. In these diseases, VEGFR2 is a well-established pharmacological target. Importantly, azidation has previously been used to tether small bioactive molecules to their receptor: light is demonstrated to trigger the covalent binding of azidated alphaxalone to the GABA receptor (*12*), leukotriene-D4 to its own receptor (*13*), p-azido-N-metilspirenon to the D2 dopamine receptors (*14*). One study even provided functional evidence that azidation can convert a reversible inhibitor into an irreversible one upon photoactivation (*15*)

As described in our first patent (*11*) by attaching the N_3_ group to a larger aromatic electron system and in a conjugated position, with no more than one σ-bond separating the N_3_ group from the delocalized electron system of a well-established inhibitor (Fig. 1), the wavelength required for photoactivation can be substantially red-shifted, as compared to the wavelength typically required for the photolysis of azides (10). As proof-of-concept molecules, here we present the derivatives (*11*) of two oxindole-based VEGFR inhibitors: sunitinib (21) and vorolanib. Both compounds exhibit absorption maxima near 450 nm, and their extended aromatic systems provide four potential conjugation sites on the indole-2-one moiety for azidation.

**Fig. 1.**
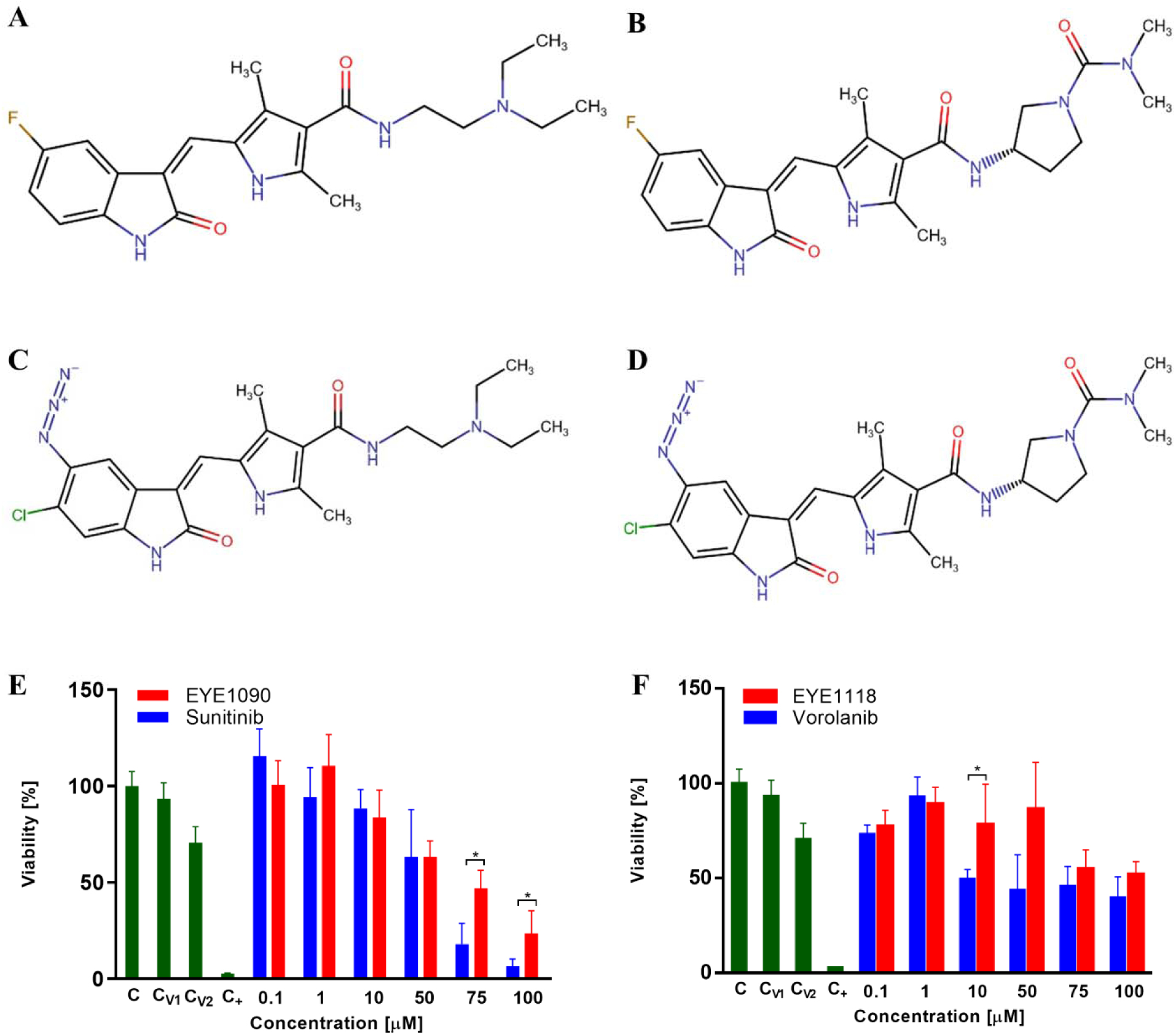
New chemical entities EYE1090 and EYE1118. **A,** Sunitinib, the parental compound of EYE1090; **B,** Vorolanib, the parental compound of EYE1118; **C,** Structure of EYE1090 (10,11); **D,** Structure of EYE1118 (10,11); **E,** Viability of NIH-3T3 cells after 12 hours incubation with sunitinib and EYE1090, “C” stands for control, “C_v1_” for 0,2% DMSO and “C_v2_” for 2% DMSO, and “C_+_” for geneticin as positive control, one asterisk indicates significance at p<0.05 level (two way ANOVA); **F,** Viability of NIH-3T3 cells after 12 hours incubation with vorolanib and EYE1118, vehicle controls statistical significance are indicated as in panel **“E”**.

Sunitinib is currently prescribed against three different types of tumors (*2*); however, its adverse effects are severe (*16*), its standard dosing regimen contains two weeks of pause (*17*), and its therapeutic range is so narrow that analytical methods had to be developed to measure its concentration in the human plasma (*18*). Light-mediated targeting may lower the required therapeutic concentration of an azidated sunitinib derivative, thereby yielding therapeutic benefit. The second scaffold, vorolanib, was previously developed to offer improved safety and faster metabolism compared to sunitinib (*19*). In spite of its less severe adverse effects and proven ocular efficiency in human subjects, the development of an oral vorolanib formulation against AMD has been stopped because of undesired hepatobiliary effects (*3,4*). This makes vorolanib a prime candidate for a modification that makes it targetable specifically to the retina by natural light seen by the patients. This approach exploits the anatomical properties of the eye, which assure that light is focused on the neuroretina, ie. right to the location where therapeutic effect needs to be exerted (*10,11*).

As described in our second patent (*10*), sunitinib azidated at the position 5 and chlorinated the position 6 of the indol-2-one moiety showed a very strong inhibition of VEGFR2 in a kinase assay with an IC_50_ value of 13 nM in the dark, which was further potentiated to 3 nM upon illumination. This molecule was termed EYE1090. Similarly, as we have previously shown (*10*), vorolanib azidated at the position 5 and chlorinated the position 6 of the indol-2-one moiety showed a very strong inhibition of VEGFR2 in a kinase assay with an IC_50_ value of 24 nM in the dark, which was further potentiated to 9 nM upon illumination. This molecule was termed EYE1118. Importantly, the chlorinated derivatives (EYE1090, EYE1118) showed superior inhibitory potential and better photoactivation properties than their counterparts lacking the halogen atom (*10*). These pharmacological properties of EYE1090 and EYE1118 favorably coincide with the „magic chlorine effect” (*20*) associated with improved gastrointestinal absorption. To assess the toxicity of the new chemical entities, we have conducted Alamar Blue assays (Fig. 1E, 1F). Interestingly, EYE1090 was less toxic than sunitinib, and this effect reached statistical significance at concentrations above 75 µM (Fig. 1E), while the same difference between EYE1118 and vorolanib reached significance at 10 µM (Fig. 1F). The solubility of the compounds prevented us from testing higher concentrations in the cytotoxicity assays. This impacted especially the comparison of vorolanib and EYE1118, which showed lower toxicity than the sunitinib scaffold and were therefore only marginally more toxic at 100 µM than the vehicle (Fig. 1E, 1F).

### Light potentiates the inhibition of overexpressed VEGFR2 by EYE1090 and EYE1118

The in vitro findings detailed above encouraged us to test the inhibition and the light effect in a cellular context. We have opted for a commercially available VEGFR2/NFAT reporter HEK293 cell line (BPS BioScience) that reports the strength of VEGF signaling as directly quantifiable luciferase activity.

To assess the light-mediated enhancement of inhibitor activity, we designed experiments involving a 10-minute-long exposure to cold white LED light (693 lux). This was intended to photolyze the azido group and trigger covalent binding of the compounds to VEGFR2, thereby increasing their inhibitory potency. Cells were cultured in 96-well plates and irradiated directly within the incubator using a Bluetooth-controlled light source, while control wells were kept in the dark.

In each experiment, we have pre-incubated the cells with the inhibitors for 50 minutes to let the molecules diffuse into the cells and bind to the intracellular portion of the VEGFR2 receptor, then we exposed the cultures to cold white LED for 10 minutes. Finally, we stimulated them with 50 µg/ml of commercially available human VEGF165 and conducted luciferase assays after having lysed the cells.

Both EYE1090 (IC_50_ = 19.06 nM, Fig. 2A) and EYE1118 (IC_50_ = 52.24 nM, Fig. 2B) showed robust inhibition of VEGFR2 signaling in the dark, yielding standard sigmoidal dose-response curves. Importantly, cold white LED illumination significantly enhanced their potency: the IC_50_ values dropped to 2.18 nM for EYE1090 and 6.49 nM for EYE1118 (p < 0.0001), indicating nearly 9-fold and 8-fold increases in inhibitory strength, respectively.

**Fig. 2.**
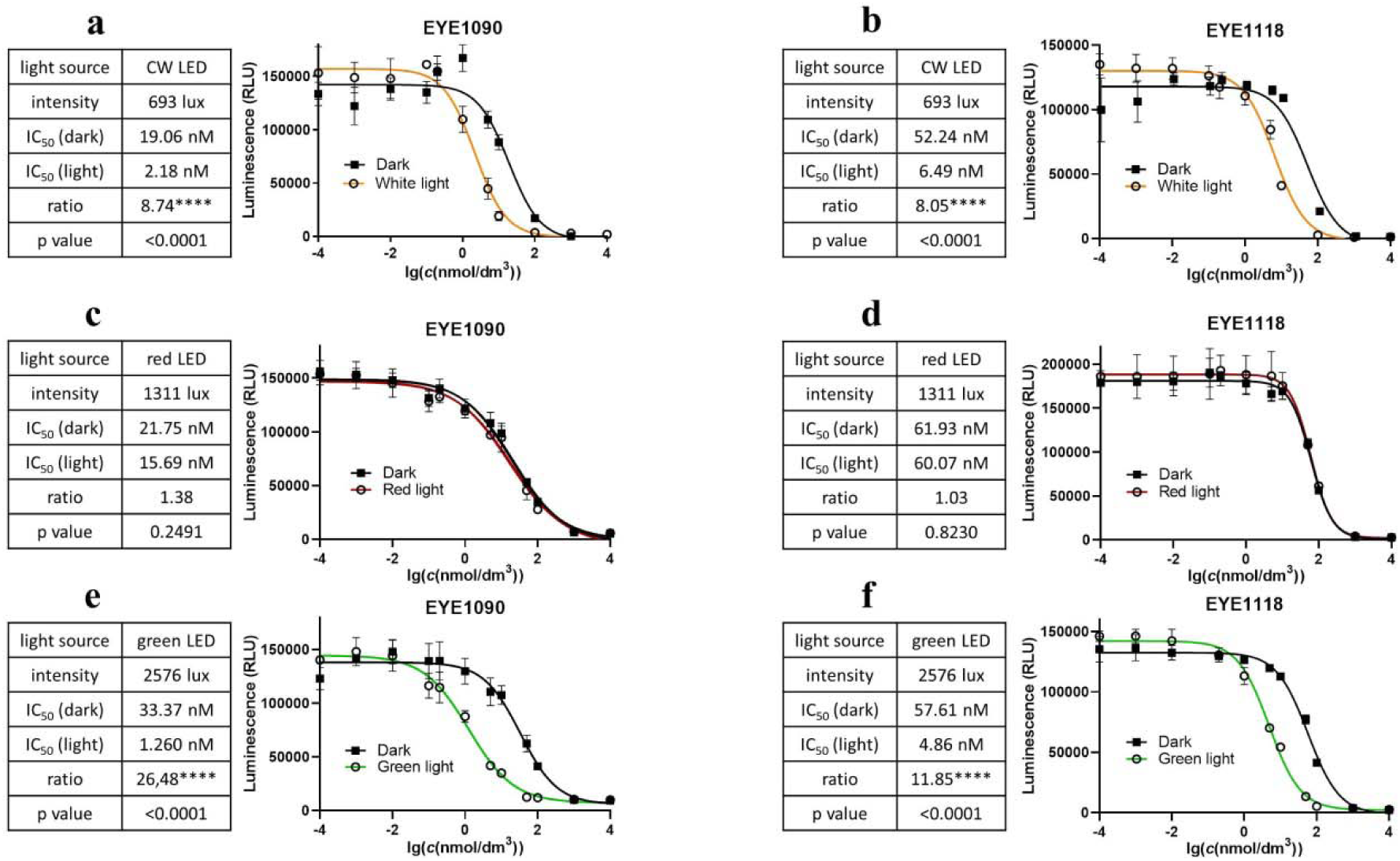
Light-dependent inhibition of the VEGF signaling in HEK cells expressing human VEGFR2 and an NFAT-luciferase construct. **a,** White light enhances 8.74-fold the inhibition by EYE1090. **b,** White light enhances 8.05-fold the inhibition by EYE1118; **c,** Red light does not enhance the inhibition by EYE1090; **d,** Red light does not enhance the inhibition by EYE1118; **e,** Green light enhances 26.48-fold the inhibition by EYE1090; **f,** Green light enhances 11.85-fold the inhibition by EYE1118. Statistical comparison of the IC_50_ values was carried out in the GraphPad Prism software.

To ensure that this enhancement was due to specific photochemical activation rather than non-specific effects of light exposure, we repeated the assay using red LED light (654 nm, 1311 lux), which has higher intensity but lower photon energy and thus cannot activate aryl azides. As expected, red light had no significant impact on the IC_50_ of either EYE1090 (Fig. 2C) or EYE1118 (Fig. 2D), supporting the specificity of the white light effect.

With an eye on medical applicability, we have finally tested a green LED with an emission peak at 516 nm. Based on its characteristics (Fig. S2), this light can effectively reach the human retina, even in older individuals or those with reduced lens transparency (*21*. Surprisingly, green light potentiated the inhibitory activity of both compounds to an even greater extent than white light: EYE1090’s IC_50_ decreased from 33.37 nM to 1.26 nM, and EYE1118’s from 57.61 nM to 4.86 nM—corresponding to a nearly 30-fold and 12-fold increases in potency, respectively (Fig. 2E, 2F; p < 0.0001 for both).

The enhanced sensitivity to green light is likely due to the expanded delocalized electron system created by the azido group, which facilitates photolysis even at wavelengths considered suboptimal for activating aryl azides. Despite a decline in the light absorption of the compounds in the green spectral range (*10*), spectral analysis of the green LED source (Fig. S1B) revealed significant emission extending into the blue region (down to ∼480 nm), allowing sufficient overlap with the compounds’ absorption spectra to initiate photolysis.

### Light changes the effect of EYE1090 and EYE1118 on VEGFR2 phosphorylation

To confirm that VEGFR2 is the molecular target mediating the light-dependent effects of our novel compounds, we have immunoblotted the receptor from the same cell line (VEGFR2/NFAT reporter HEK293) that we used to unravel the functional effect (Fig. 2) of the signaling pathway. We first characterized the kinetics of VEGFR2 phosphorylation in response to VEGF stimulation (Fig. S2), then proceeded to assess the impact of EYE1090 and EYE1118 on receptor phosphorylation after 10 minutes of VEGF treatment and under controlled lighting conditions.

Cells were pre-treated with either EYE1090, EYE1118, or vehicle control (DMSO) under dim red light to prevent unintended photoactivation. Cultures were then exposed to the same white LED light as used in luciferase experiments or kept in the dark. Following irradiation, cells were stimulated with 50 ng/mL of VEGF165 and harvested 10 minutes later. Lysates were analyzed by SDS-PAGE and Western blotting to assess the phosphorylation status of two key tyrosine residues: Y951 and Y1175.

Tyrosine 951 is implicated in endothelial cell survival and vascular permeability (*22*), and is required for the subsequent phosphorylation of additional tyrosines on the intracellular domain of VEGFR2 (*23*). Tyrosine 1175 is critical for multiple endothelial functions, including permeability, proliferation, and migration (*22*), and it activates several downstream effectors, including protein kinase C (PKC), focal adhesion kinase (FAK), phosphoinositide 3-kinase (PI3K), and p38 mitogen-activated protein kinase (p38 MAPK)(22).

As expected, VEGF165 stimulation induced robust phosphorylation of both Y951 and Y1175 (Fig. 3). In the absence of light, neither EYE1090 nor EYE1118 substantially inhibited this phosphorylation, even at 100 nM. However, when combined with light exposure, both compounds significantly suppressed phosphorylation. At 100 nM, EYE1090 and EYE1118 reduced phosphorylation levels to baseline (comparable to unstimulated controls), while 50 nM of either compound produced a statistically significant reduction relative to VEGF alone.

**Fig. 3.**
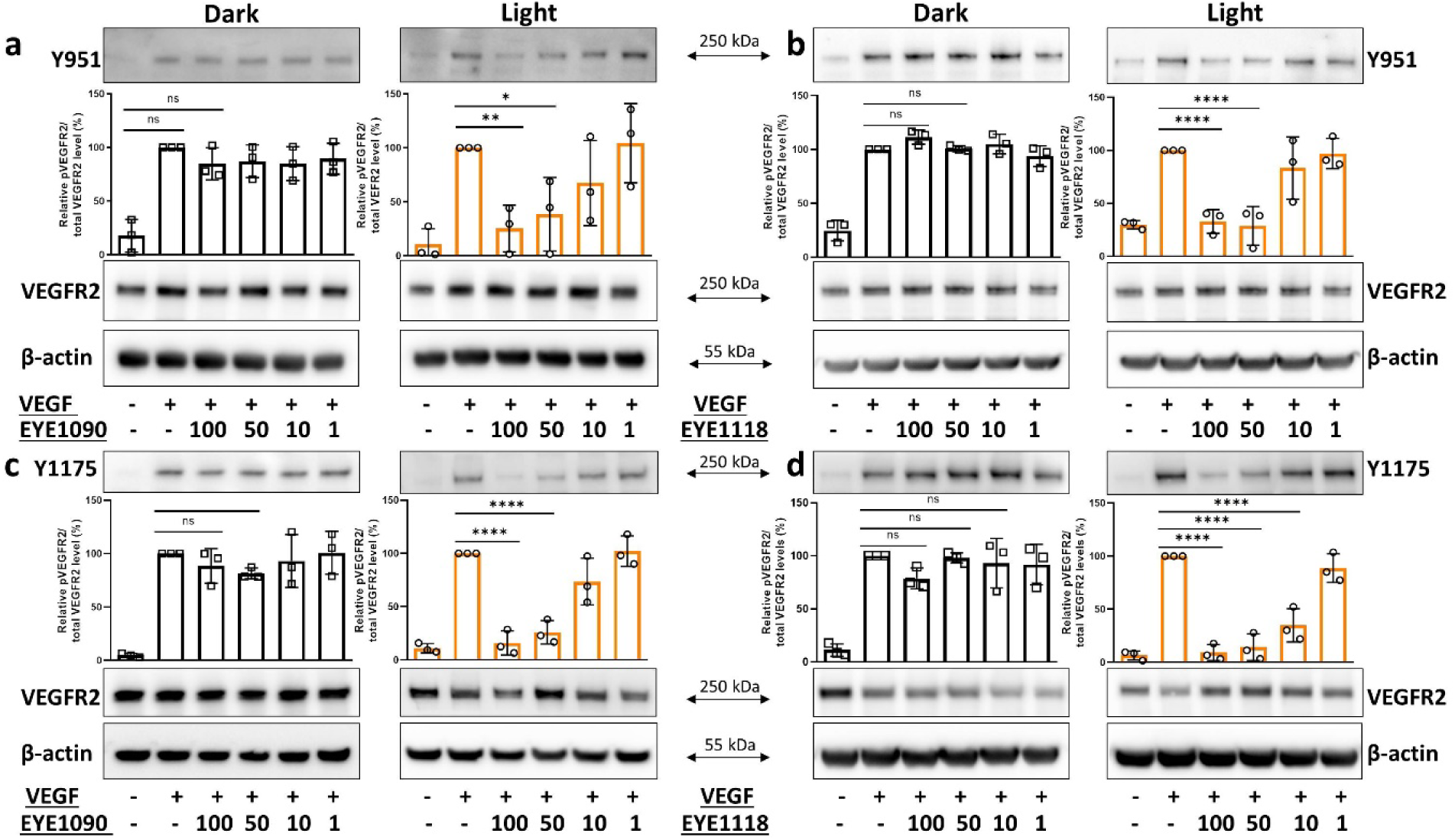
Light-dependent effect of EYE1090 and EYE1118 on VEGF-induced VEGFR2 phosphorylation. For each panel, the upper blots show the amount of VEGFR2 phosphorylated at Y951 (**a, b**) and at Y1175 (**c, d**). The middle blots show the total amount of VEGFR2, the lower blots show the amount of β-actin. The quantifications are based on three independent experiments identical to those shown in the blots. Statistical analysis was carried out in the GraphPad Prism software. The concentrations of EYE1090 (**a, c**) and EYE1118 (**b, d**) are indicated in nM units.

To confirm that these effects were not due to altered protein levels, membranes were stripped and reprobed with antibodies against β-actin and total (including non-phosphorylated) VEGFR2. β-actin served as a loading control (lower panels, Fig. 3A–3D), confirming consistent sample input, while total VEGFR2 levels remained unchanged (middle panels, Fig. 3A–3D), indicating that the reduced phosphorylation was not due to degradation or downregulation of the receptor. These results strongly support our core hypothesis that light exposure triggers covalent anchoring of EYE1090 and EYE1118 to VEGFR2, thereby preventing its activation.

A similar light-dependent effect was observed for EYE1118 at tyrosine 1214 (Fig. S3), which contributes to the reorganization of the actin cytoskeleton. However, due to high basal phosphorylation of Y1214 in the absence of VEGF, quantification of this effect was not feasible.

### Light-dependent inhibition of the angiogenic and migratory potential of endothelial cells

Microvascular endothelial cells are widely used to test the pro- and anti-angiogenic potential of drug candidate molecules. This is attributable to their capability of spontaneously forming a network of tubular structures on extracellular matrices. Such structures share many features with real capillaries. The emergence of the network is enabled by the presence of all the required receptors and actuator proteins in the endothelial cells (*24*).

Given the intended therapeutic application of EYE1090 and EYE1118 in ocular conditions, relying on natural light reaching the retina, we selected primary human retinal microvascular endothelial cells (HRMECs) derived from post-mortem donors (ACBRI-181, Cell Systems) to assess the anti-angiogenic effects of the compounds. These cells readily form complex tubular networks on Geltrex (Thermo Fisher), exhibiting multiple quantifiable angiogenic features even in the absence of exogenous VEGF (*25*).

Based on a preliminary experiment (Supplementary Video), HRMECs were incubated with EYE1090 or EYE1118 for 12 hours, then loaded with calcein, and imaged (representative networks in Fig. 4A and 5A). We quantified two parameters with a custom-developed algorithm (Fig S4) in the Nikon NIS-Elements platform: the number of closed loops (“meshes”) (Fig. 4B and 5B), and the number of nodes (Fig. 4C and 5C). Both EYE1090 and EYE1118 exhibited robust inhibition of HRMEC tube formation in the dark.

**Fig. 4.**
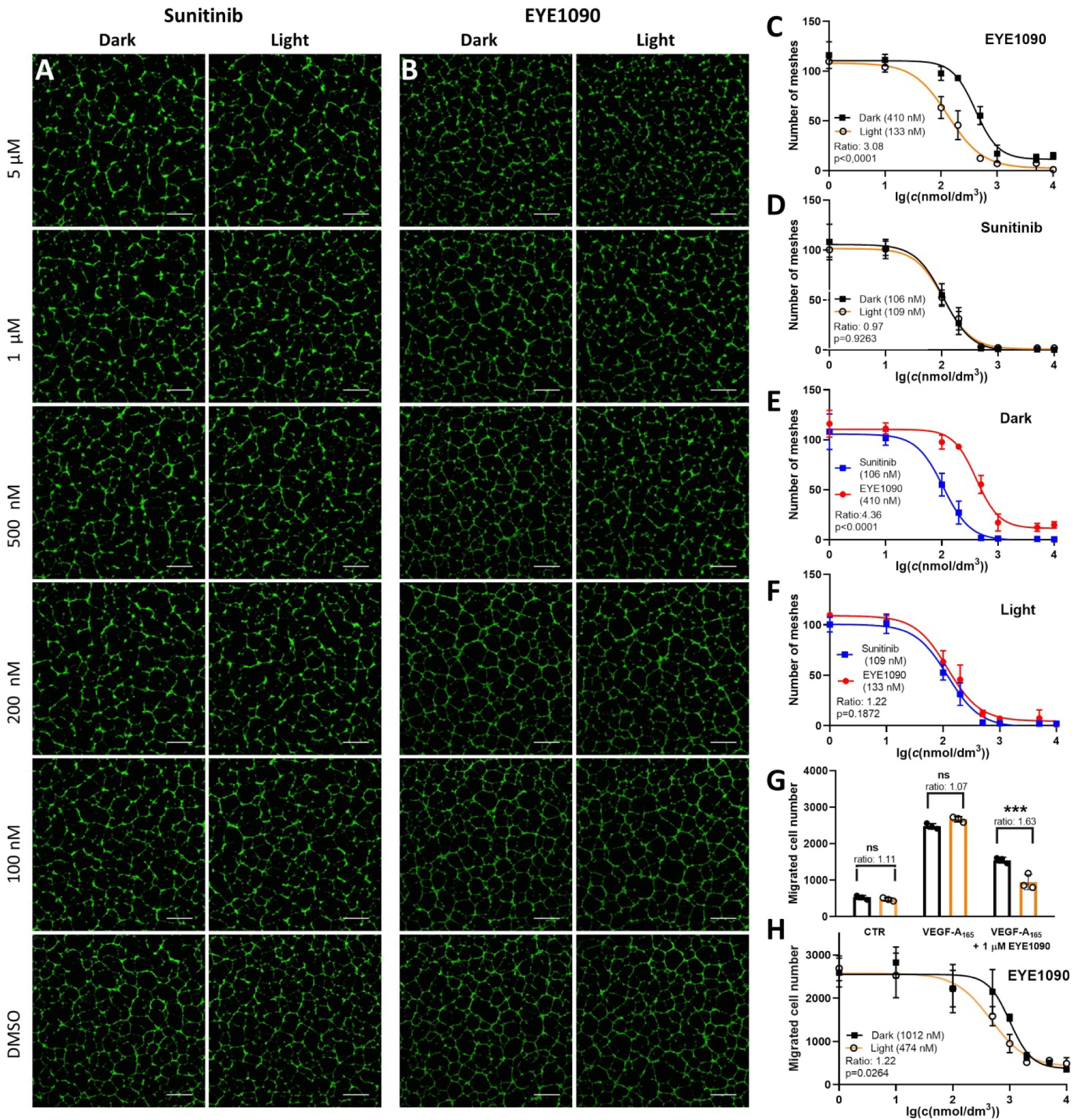
The light-dependent inhibition of the angiogenic potential and migration of HRMEC by EYE1090. **A,** Networks emerging on a Geltrex matrix showing the inhibition by sunitinib that does not change with light along with the vehicle control at the bottom of the panel, scalebar: 250 µm; **B,** Networks emerging on a Geltrex matrix showing the inhibition by EYE1090 which is potentiated by light (right column), vehicle control is shown at the bottom of the panel, scalebar: 250 µm; **C,** Comparison of the number of the meshes and IC_50_ values of EYE1090 between light and dark conditions; **D,** Comparison of the number of the meshes and IC_50_ values of sunitinib between light and dark conditions; **E,** Comparison of the number of the meshes and IC_50_ values between EYE1090 and sunitinib, without pre-irradiation; **F,** Comparison of the number of the meshes and IC_50_ values between EYE1090 and sunitinib, with 10 minutes of pre-irradiation before tube formation; **G,** Light significantly potentiates the inhibiton of HRMEC migration by EYE1090, while no light effect is observed on cells treated with vehicle alone or VEGF and vehicle; **H,** The IC_50_ value of EYE1090 based on the number of migrated cells decrease significantly upon illumination. For a direct statistical comparison of the IC_50_ values, the built-in option in GraphPad Prism software was used for panels “C”, “D”, “E”, “F”, and “H”.

**Fig. 5.**
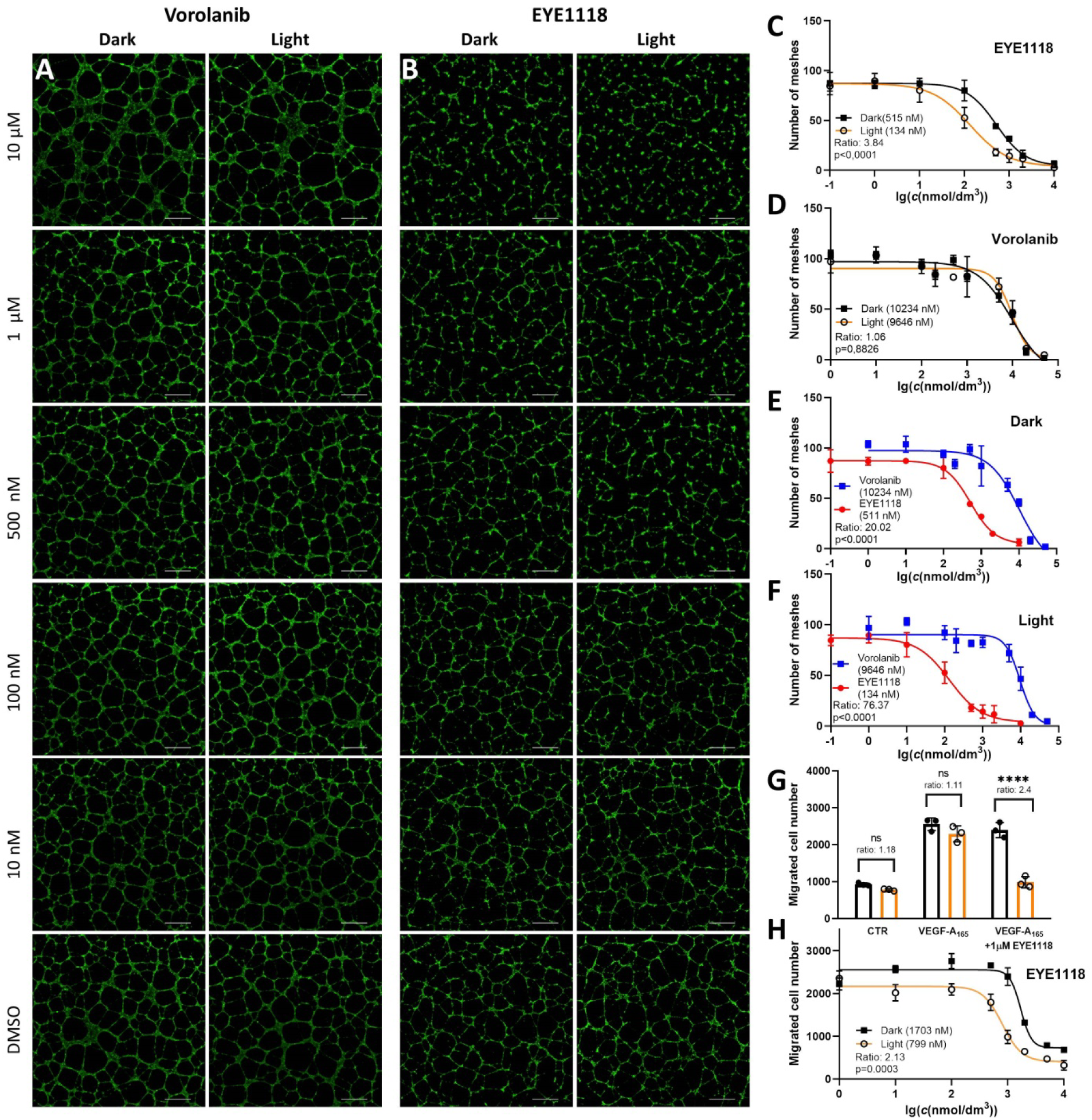
The light-dependent inhibition of the angiogenic potential and migration of HRMEC by EYE1118. **A,** Networks emerging on a Geltrex matrix showing the inhibition by vorolanib that does not change with light along with the vehicle control at the bottom of the panel, scalebar: 250 µm; **B,** Networks emerging on a Geltrex matrix showing the inhibition by EYE1118 which is potentiated by light (right column), vehicle control is shown at the bottom of the panel, scalebar: 250 µm; **C,** Comparison of the number of the meshes and IC_50_ values of EYE1118 between light and dark conditions; **D,** Comparison of the number of the meshes and IC_50_ values of vorolanib between light and dark conditions; **E,** Comparison of the number of the meshes and IC_50_ values between EYE1118 and vorolanib, without pre-irradiation; **F,** Comparison of the number of the meshes and IC_50_ values between EYE1118 and vorolanib, with 10 minutes of pre-irradiation before tube formation; **G,** Light significantly potentiates the inhibiton of HRMEC migration by EYE1118, while no light effect is observed on cells treated with vehicle alone or VEGF and vehicle; **H,** The IC_50_ value of EYE1118 based on the number of migrated cells decrease significantly upon illumination. For a direct statistical comparison of the IC_50_ values, the built-in option in GraphPad Prism software was used for panels “C”, “D”, “E”, “F”, and “H”.

To investigate the effect of light, HRMECs were preincubated with the compounds, then briefly irradiated with cold white LED light (693 lux) prior to seeding onto Geltrex, following the same assay protocol. Despite the reasonable expectation that a substantial proportion of the azidated inhibitors would be photolyzed during the short illumination, light exposure still led to significantly stronger inhibition of tube formation for EYE1090 (Fig. 4A, 4C) compared to the dark condition, where the compound was present throughout the entire 12-hour incubation. This likely reflects the covalent binding of a sufficient fraction of photoactivated inhibitor to VEGFR2 at the time of light exposure.

To compare EYE1090 to its parental compound, sunitinib, we have conducted tubulogenesis assays with the latter within the same conditions. We have found, in line with the chemical core concept, that the efficiency of sunitinib did not change upon irradiation (Fig. 4B, 4D). While in the dark, sunitinib inhibited tubulogenesis slightly more efficiently than EYE1090 (Fig. 4E), this difference vanished after 10 minutes of pre-exposure to light when the novel compound became non-inferior in terms of inhibitory potential (Fig. 4F). Of note, this short illumination before a 12-hour-long experiment was sufficient to equalize the efficiency of EYE1090 and sunitinib in spite of the obviously occurring VEGFR2 re-synthesis and the presumed ectopic photoactivation of EYE1090 resulting in binding to non-target biomolecules.

Initially, we did not observe a light-potentiated effect of EYE1118 on angiogenesis. Since its parent compound, vorolanib, is known to interact with the efflux transporter ABCG2 (*26*), we hypothesized that a significant portion of EYE1118 was being actively exported from the cells. To test this, we included the ABCG2-specific inhibitor Ko-143 in all experiments with EYE1118. Under these conditions, a clear and statistically significant light-enhanced inhibition of angiogenesis emerged (Fig. 5A, 5C). Notably, Ko-143 also improved the inhibitory effect of EYE1118 in the dark (Fig. S5A–S5C), suggesting that ABCG2-mediated efflux reduces intracellular drug levels throughout the assay duration.

As we did for EYE1090, we compared EYE1118 to its parental compound, vorolanib. We added Ko-143 to all of these endothelial cultures, given the reported interaction with ABCG2 (*26*). Expectedly, we have found that light did not change the IC_50_ value of vorolanib (Fig. 5B, 5D). A striking difference emerged between EYE1118 and vorolanib in the dark (Fig. 5E), the former being approximately a 100-fold stronger inhibitor of tubulogenesis. The superior efficiency of the novel compound manifested even more in pre-irradiated cultures (Fig. 5F). The difference observed in these experiments exceeded that in biochemical assays already favoring EYE1118 over vorolanib (Fig. 1F), which may be attributed to the different trafficking or subcellular localization of the two tyrosine-kinase inhibitors in microvascular endothelial cells from human retinae.

Together, these results demonstrate that both EYE1090 and EYE1118 effectively suppress *in vitro* angiogenesis in HRMECs and exhibit markedly increased inhibitory potential under light exposure. These effects are evident both in quantitative measurements (number of meshes; Fig. 4C–4F and 5C–5F) and in the qualitative morphology of the tubular networks (Fig. 4A-4B and 5A-5B), underscoring their promising anti-angiogenic potential upon illumination. Moreover, unexpectedly, EYE1118 represses tubulogenesis in the dark much more potently than vorolanib (Fig. 5F), its parental compound already tested against AMD in human patients.

We further investigated the effect of the compounds on HRMEC migration using a transwell assay format. Cells were serum-starved overnight, preincubated with inhibitors, then seeded onto the upper chamber of transwell inserts. Following a 10-minute white light exposure, cells were allowed to migrate for 6 hours toward VEGF165 placed in the lower chamber as a chemoattractant. Migrated cells were quantified by nuclear staining (Fig. S6). As with the tube formation assay, Ko-143 was included in all EYE1118 conditions to block ABCG2 activity.

Both EYE1090 and EYE1118 effectively inhibited HRMEC migration in the dark, with IC_50_ values of 1032 nM and 1706 nM, respectively (Fig. 4G, 4H, 5G, 5H). Light exposure significantly enhanced the inhibitory effect: IC_50_ values decreased to 484 nM for EYE1090 (Fig. 4H) and to 756 nM for EYE1118 (Fig. 5H), confirming that photoactivation potentiates the compounds’ ability to disrupt VEGFR2-mediated migration. These results align with our findings on VEGFR2 Y1175 phosphorylation, a key site governing endothelial motility.

In summary, our *in vitro* data show that both EYE1090 and EYE1118 exert broad VEGFR2-inhibitory effects across multiple cellular functions, including angiogenesis and migration. The enhancement of these effects by light across distinct functional readouts supports the notion that photoactivation strengthens the association of these compounds with VEGF receptors in live cells. The results provide compelling evidence for the feasibility of light-targeted anti-angiogenic therapies.

### Light influences the biodistribution of EYE1090 in vivo in albino mice

Encouraged by the promising light-dependent effects observed *in vitro,* we conducted a small-scale pharmacokinetic study in BALB/c mice, which lack ocular melanin and are therefore not expected to sequester sunitinib-like compounds in the uvea (*27*). In these experiments, animals received EYE1090 via oral gavage under dark conditions. Thirty minutes post-administration, one group was exposed to white light at 1000 lux, while the control group remained in dim red light. Four hours after gavage, all animals were sacrificed under dim red light to prevent unintended photoactivation.

Serum levels of EYE1090 were quantified using a liquid–liquid extraction method adapted from established protocols for sunitinib (*17*). Notably, white light exposure resulted in a reduction of more than 50% in the mean serum concentrations of EYE1090 compared to dark-treated controls. Moreover, a statistically significant increase in serum levels was observed only in the dark condition, whereas light-exposed animals did not show a significant change relative to baseline (Fig. 6).

**Fig. 6.**
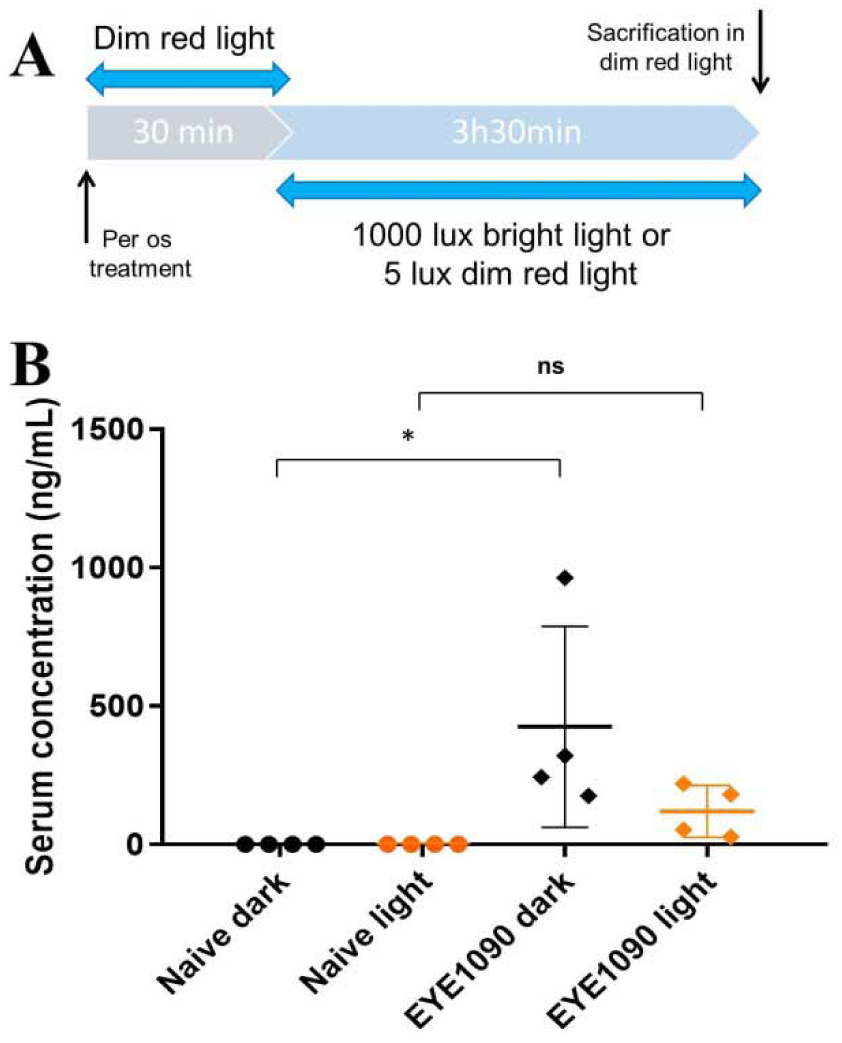
Amount of EYE1090 in the mouse serum as measured by liquid-liquid extraction followed by HPLC. **A,** Experimental design; **B,** EYE1090 serum concentrations increase significantly (p = 0.0331) 4 hours after EYE1090 treatment, when the animals are kept in the dark, while do not increase significantly (p = 0.8051) in animals kept in the light. Results of a one-way ANOVA are shown as calculated by the Graphpad Prism software.

In these experiments, we cannot exclude the possibility of EYE1090 photoactivation in non-ocular tissues including superficial blood vessels located in thinner regions of the skin. Moreover, large standard deviations of the serum concentrations were observed, probably due to individual differences in EYE1090 metabolism. Nevertheless, these results support the hypothesis that EYE1090 undergoes photoactivation in the retina, which, by anatomical design, captures ambient light with outstanding efficiency.

These findings prompted us to evaluate the in vivo efficacy of EYE1090 and EYE1118 in models of pathological ocular neovascularization. Specifically, we tested their effects on VEGF-induced vascular permeability following intravitreal injection in Long Evans rats, as well as their therapeutic impact in a laser-induced choroidal neovascularization (CNV) model in C57BL/6 mice. These pigmented strains were deliberately chosen to account for potential uveal accumulation of EYE1090, given their melanin content.

### EYE1090 and EYE1118 in decreases VEGF-induced retinal leakage in rats

One of the principal targeted medical indications for our engineered molecules is the group of retinopathies characterized by neovascularization and/or severe edema. In these conditions, the anatomical structure of the eye, specifically its natural ability to focus incident light onto the retina, can be strategically exploited to locally activate EYE1090 and EYE1118. This allows light-mediated binding of the compounds to VEGFR2, a key driver of pathological angiogenesis in such diseases. This targeting strategy, however, imposes constraints on the required activation wavelength and limits the pool of kinase inhibitors with appropriate chemical scaffolds that can be azidated and photoactivated within the ocular environment.

Among the two candidates, EYE1118 is particularly well-suited for ocular applications. It offers a more favorable systemic safety profile (*19*), exhibits minimal ocular side effects (*28*), and is derived from vorolanib which is a compound already tested in human clinical trials for age-related macular degeneration (AMD), where it demonstrated strong efficacy (*3,4*). We envision that light-directed retinal activation of EYE1118 will allow for a substantial dose reduction compared to the systemic doses used in previous oral vorolanib trials for AMD.

Despite the higher likelihood of systemic adverse effects due to its sunitinib-based scaffold, we also included EYE1090 in our *in vivo* studies. This decision was supported by multiple factors: sunitinib has already been evaluated in ocular applications via intravitreal administration (*29*), and it is known to accumulate significantly in uveal tissues due to melanin binding (*30*). EYE1090 is expected to exhibit similar melanin-binding properties, potentially forming a depot within the uvea. Given that this depot may be shielded from incident light by the retinal pigment epithelium (RPE), including EYE1090 in our comparison offered an opportunity to assess the impact of pigment-related sequestration relative to EYE1118, which is predicted to bind melanin to a lesser extent (*27*).

To assess retinal vascular permeability, we employed the Evans Blue assay (*31*) and relied on the fluorescence-based detection of this azo dye (*32*) (Fig. S7). Our model involved intravitreal injection of VEGF165, a well-established method to provoke capillary leakage via VEGFR2 activation (*25*), inducing permeability changes comparable to those seen in streptozotocin-induced diabetic retinopathy (*33*). We used a VEGF165 dose of 50 ng/eye, which in our lab consistently produces a more pronounced increase in retinal permeability than diabetic models.

To evaluate the therapeutic potential of our compounds in this model, EYE1090 and EYE1118 were administered orally at 1 mg/kg daily for three days prior to the intravitreal VEGF injection (Fig. 7A). Notably, this dosage corresponds to a human equivalent dose (HED) of approximately 0.16 mg/kg, which is substantially lower than doses previously used in animal (*34*) or human (*4*) studies involving vorolanib. In this context, our engineered compounds demonstrated approximately tenfold higher efficacy than previously tested tyrosine kinase inhibitors for ocular indications. This highlights their potential as a new class of highly potent, retina-targeted therapeutics (Figure 7C).

**Fig. 7.**
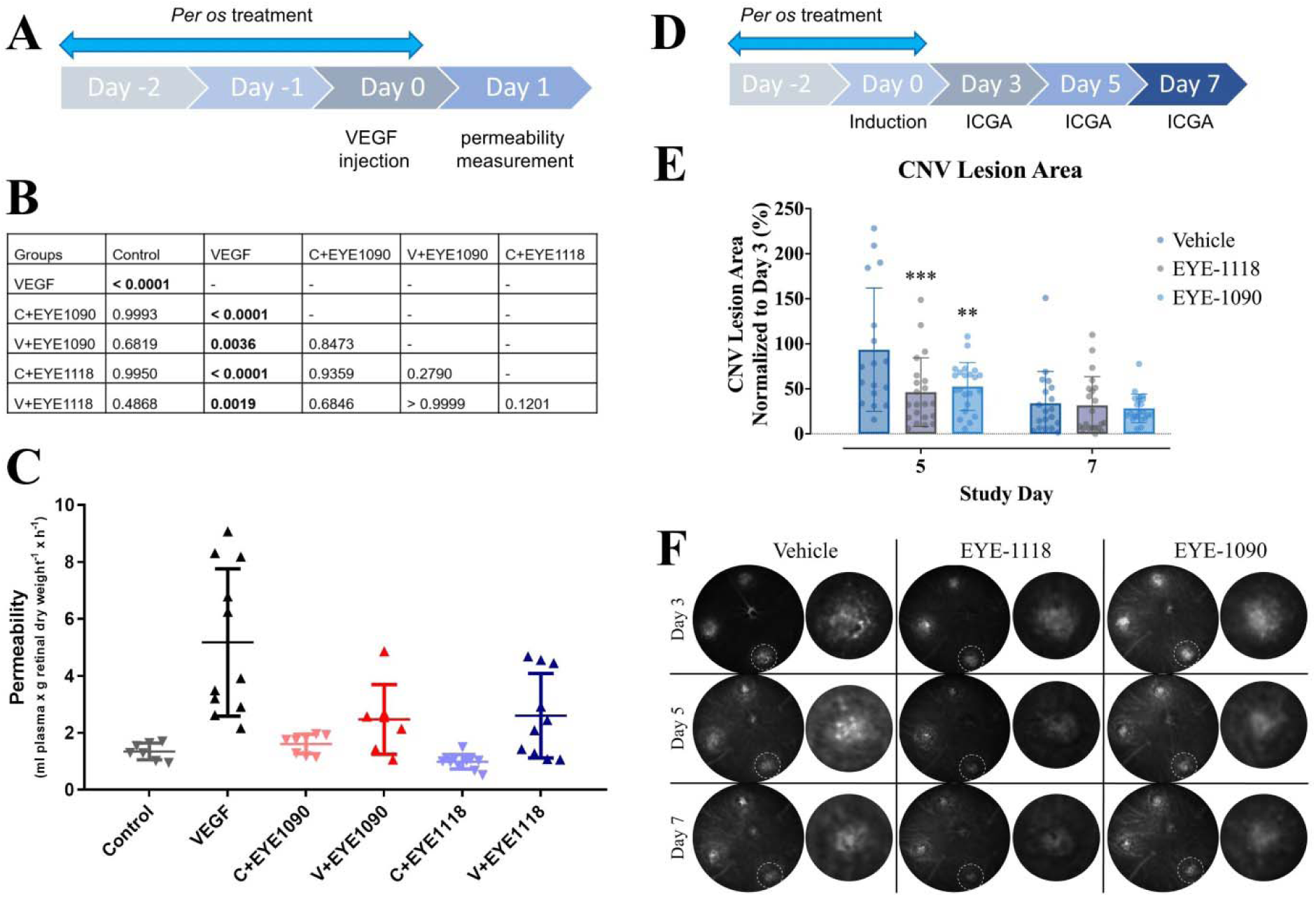
The efficacy of orally administered EYE1090 and EYE1118 in rodent models of ocular vasculopathies. **A,** The experimental design for testing the effect on retinal permeability in Long Evans rats. Animals were treated for three consecutive days with 1 mg/kg of inhibitors or vehicle. VEGF was injected intravitreally into the left eye of each animal, while the right eye received sterile salt solution in an equivalent amount. Permeability measurements were performed 24 hours later; **B,** Statistical analysis of permeability values from Long Evans rats. Data were analyzed by one-way ANOVA and Tukey’s post-hoc test in GraphPad Prism; **C,** Graphical representation of the permeability values from Long Evans rats. For animals treated with compounds, “C” stands for control eyes and “V” for VEGF-injected eyes; **D,** The experimental design for testing the efficiency of prophylactically administered 1 mg/kg EYE1090 and 1 mg/kg EYE1118 in a C57/BL6 mouse model of choroidal neovascularization; **E,** Quantification of the lesion area, 3 asterisks meaning p<0.001 (EYE1118); 2 asterisks meaning p<0.01 (EYE1090). Lesion area on day 3 is set as the reference value (100%); **F,** Example photos of laser-generated lesions at days 3, 5, and 7 for each treatment group. Insets show the magnification of a single CNV spot.

### EYE1090 and EYE1118 decrease the lesion area in a CNV model

Encouraged by the efficacy of EYE1090 and EYE1118 in the rat model, we proceeded to evaluate their therapeutic potential in a mouse model of CNV (*35*) which is a defining pathological feature of wet AMD. We decided to administer the compounds in a prophylactic manner to uncover whether they can attenuate the burden of the retinal damage triggered at a later time point.

Male C57BL/6 mice received daily oral gavage of either EYE1090 or EYE1118 at a low dose of 1 mg/kg for three consecutive days (Fig. 7D), while control animals received vehicle only. On the third day, we induced CNV by disrupting the Bruch’s membrane at three predefined sites in the fundus using a targeted laser, allowing choroidal vessels to invade the subretinal space. This procedure closely mimicks the neovascular pathology observed in wet AMD (*35*). Lesions were monitored on days 3, 5, and 7 post-laser treatment using indocyanine green angiography (Fig. 7F), which was selected to avoid spectral interference with the intrinsic fluorescence of our oxindole-based compounds.

Consistent with previous reports of spontaneous CNV regression over time (*36*), we observed a general reduction in lesion size by day 7. However, already by day 5, animals pretreated with EYE1118 exhibited a very highly significant reduction in lesion area, while EYE1090 treatment led to a highly significant decrease compared to controls (Fig. 7E).

Taken together, these findings provide compelling evidence that EYE1090 and EYE1118 not only inhibit pathological vascular responses when administered prophylactically but may also limit the extent of choroidal neovascularization and preserve retinal integrity in AMD. These results strongly support further investigation of these compounds as light-targetable, systemically administered therapeutics for the prevention and early intervention of neovascular retinal diseases.

### Preliminary assessment of hepatotoxicity

A major hurdle in the clinical development of our compounds is the potential for hepatobiliary toxicity. Although sunitinib is an approved anticancer agent, its clinical use is constrained by the risk of serious, and in some cases life-threatening, hepatotoxicity (*37*). Numerous reports have documented cases of sunitinib-induced liver failure in patients (38). In clinical trials, 2–5% of patients treated with sunitinib experienced grade 3 or 4 elevations in liver enzymes (*39*), and hepatic failure occurred in approximately 0.3% of cases (*40*). Similarly, for the second scaffold used in this study, namely vorolanib, severe hepatotoxicity was the primary reason for halting its development as an oral therapy for AMD (*4*).

Our chemical modification was designed to mitigate these risks by reducing systemic exposure through tissue-specific activation of the drug. In addition to this novel, light-based targeting strategy, the chemical modification of the oxindole scaffold at position 5 itself may confer a further safety advantage. In sunitinib, this position is occupied by a fluorine atom, which is known to undergo oxidative defluorination, producing a reactive quinoneimine intermediate implicated in metabolic toxicity (*40*). This bioactivation pathway is believed to be central to sunitinib’s hepatotoxic profile. Replacing fluorine with an azido group, as in EYE1090 and EYE1118, could potentially block this metabolic route and reduce hepatotoxic risk.

To evaluate the hepatotoxicity of our new compounds, we performed an acute toxicity study in BALB/c mice. Animals received a single oral dose of 40 mg/kg of either EYE1090 or EYE1118 and were maintained under either standard light/dark cycles (12 h/12 h) or continuous dim red light. Three days post-administration, livers were harvested for histopathological examination. Hematoxylin and eosin staining was used to assess liver architecture, including the integrity of portal tracts, periportal hepatocytes, sinusoidal organization, and centrilobular zones (Figure 8A, 8B, upper panels). Across all conditions, these structures remained intact, with no evidence of necrosis or overt cellular stress.

**Fig. 8.**
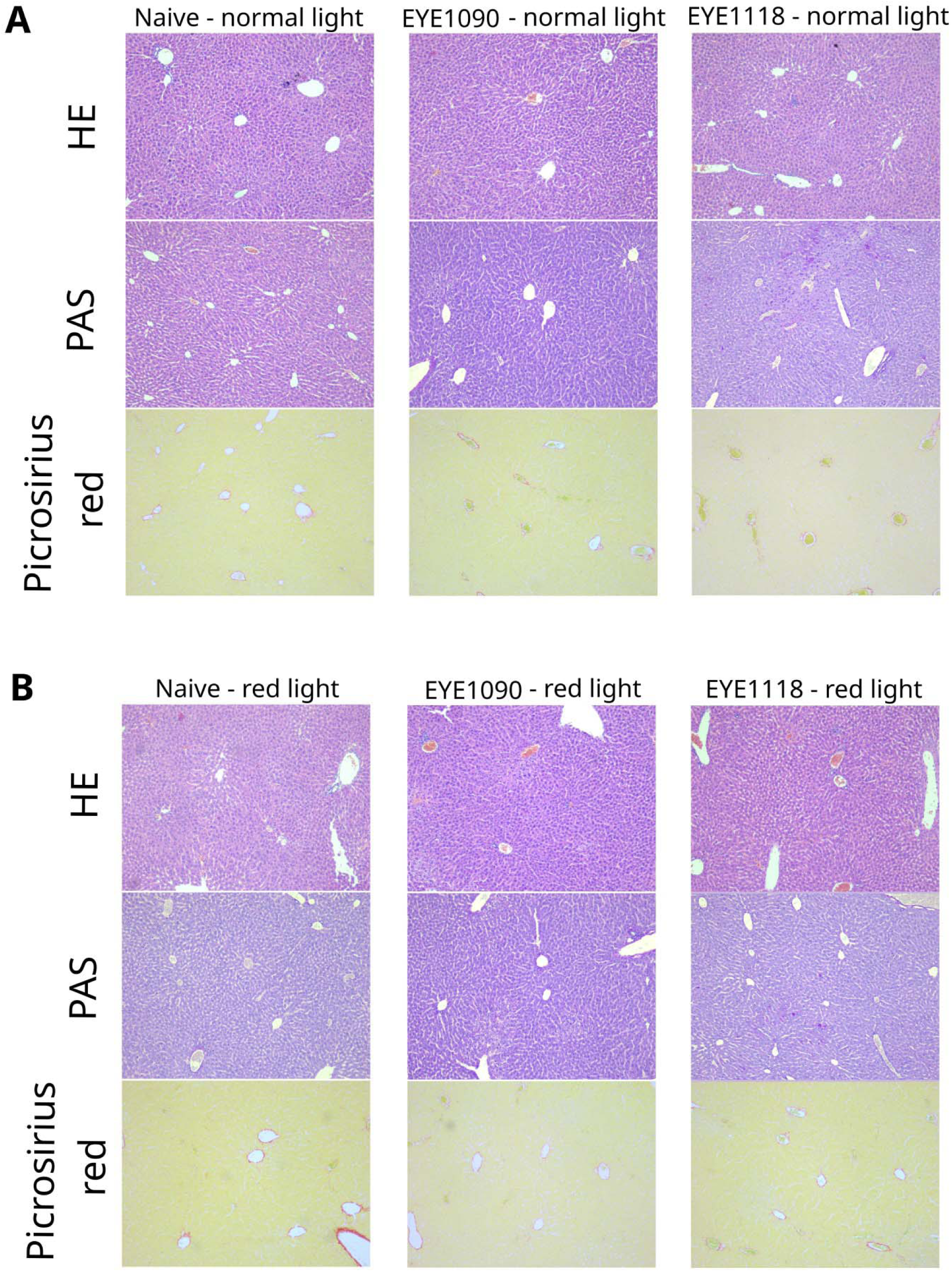
Hepatotoxicity assessment of EYE1090 and EYE1118 in BALB/c mice. **A,** Hematoxylin-eosin, PAS, and Picrosirius Red staining of liver samples from animals kept in normal light; **B,** Hematoxylin-eosin, PAS, and Picrosirius Red staining of liver samples from animals kept dim red light.

Periodic acid–Schiff (PAS) staining confirmed normal glycogen storage in hepatocytes, and no increase in Kupffer cell number was observed (Fig. 8A, 8B, middle panels). Additionally, picrosirius red staining revealed no signs of fibrosis or abnormal collagen deposition (Fig. 8A, 8B, lower panels).

## DISCUSSION

Previous clinical investigations have shown that although orally administered vorolanib is highly effective against AMD, its therapeutic potential is compromised by hepatobiliary toxicity, which ultimately halted its development for ophthalmic indications (*3, 4*).

In this study, to overcome these difficulties, we introduce a fundamentally new strategy for achieving strong, potentially irreversible receptor inhibition through targeted photoactivation of azidated receptor ligands. This approach, which we coin *optotargeting,* leverages azido-functionalized inhibitors that retain binding affinity in the dark and can be converted into covalent binders by illuminating the target tissue. This concept imposes specific chemical design constraints: the parent molecule must absorb visible light, and the azido group must be positioned in conjugation with the molecule’s delocalized π-electron system. Our previous results demonstrate that azidation itself can enhance the inhibitory activity of oxindole-based scaffolds (*10*), even prior to illumination. Moreover, the general principles outlined here may apply to a broader range of drug classes, including unrelated agonists and antagonists.

Critically, this light-based targeting strategy offers a unique opportunity in ophthalmology. Since the eye’s optical system is anatomically and functionally optimized to focus incoming light onto the retina, natural light exposure is expected to selectively photoactivate compounds like EYE1118 precisely where their pharmacological effect is needed. This mechanism would allow for retina-specific activation of the drug, reducing systemic exposure and enabling lower therapeutic dosing.

Supporting this, both EYE1090 and EYE1118 significantly reduced VEGF-induced retinal vascular leakage in a rat model, even when challenged with a dose of VEGF (50 µg) that exceeds typical pathological levels seen in diabetic retinopathy (*41, 42*) (Fig. 7C). Strikingly, prophylactic administration of these compounds at a low dose of 1 mg/kg also reduced lesion size in a mouse model of CNV (Fig. 7E, 7F), suggesting robust preventive efficacy. Additionally, our acute toxicity studies in mice revealed no signs of hepatotoxicity even at a forty-fold higher dose (Fig. 8), highlighting the favorable safety profile of both compounds at therapeutically relevant concentrations.

EYE1090, derived from sunitinib, may present greater systemic toxicity risks, but also holds promise. The known melanin-binding properties of sunitinib suggest that EYE1090 may accumulate in pigmented ocular tissues such as the uvea, potentially creating a depot effect.

Depending on the pharmacokinetics and light exposure regimen, this could be either a limitation or an opportunity for sustained local delivery. Moreover, the oncological potential of EYE1090 is substantial. The compound could be activated with light in solid tumors or metastatic sites during surgical exposure, or through endoscopic illumination of tumors in accessible anatomical cavities.

In all these contexts, targeted light exposure would reduce the systemic dose needed to achieve therapeutic efficacy, thereby lowering the risk of toxicity compared to conventional systemic sunitinib administration (*17*).

In summary, our results provide strong proof-of-concept that azidated kinase inhibitors can be selectively activated by light to exert enhanced and potentially irreversible effects at the intended site of action. EYE1118 is a promising candidate for treating retinal vascular diseases with the potential to combine high efficacy, tissue specificity, and improved safety through light-guided pharmacology.

## Supporting information

Video showing angiogenesis from HRMEC

Spectra of the light sources use in the study

Kinetics of VEGFR2 phosphorylation upon addition of VEGF

Effect of EYE1118 on VEGFR2 phosphorylation at Y1214

Methodology for quantifying in vitro angiogenesis from HRMEC

Effect of ABCG2 inhibition on the efficiency of EYE1118

Methodology for quantifying migration of HRMEC

Calibration curve for Evans Blue measurement in rats

## Acknowledgments

We do thank for the technical assistance from Viktória Szabóné Honfi.

## Author contributions

KAK designed research, BB, RA, LT, BL, NJ, ASz, ÁL, APK, AB performed research, BB, MCG, ÁL, AB analyzed data, KAK wrote the paper.

