## Supplementary figures and images for "Three nitrogen atoms turn old kinase inhibitors into new targetable remedy"

### Calibration curve for Evans Blue measurement in rats

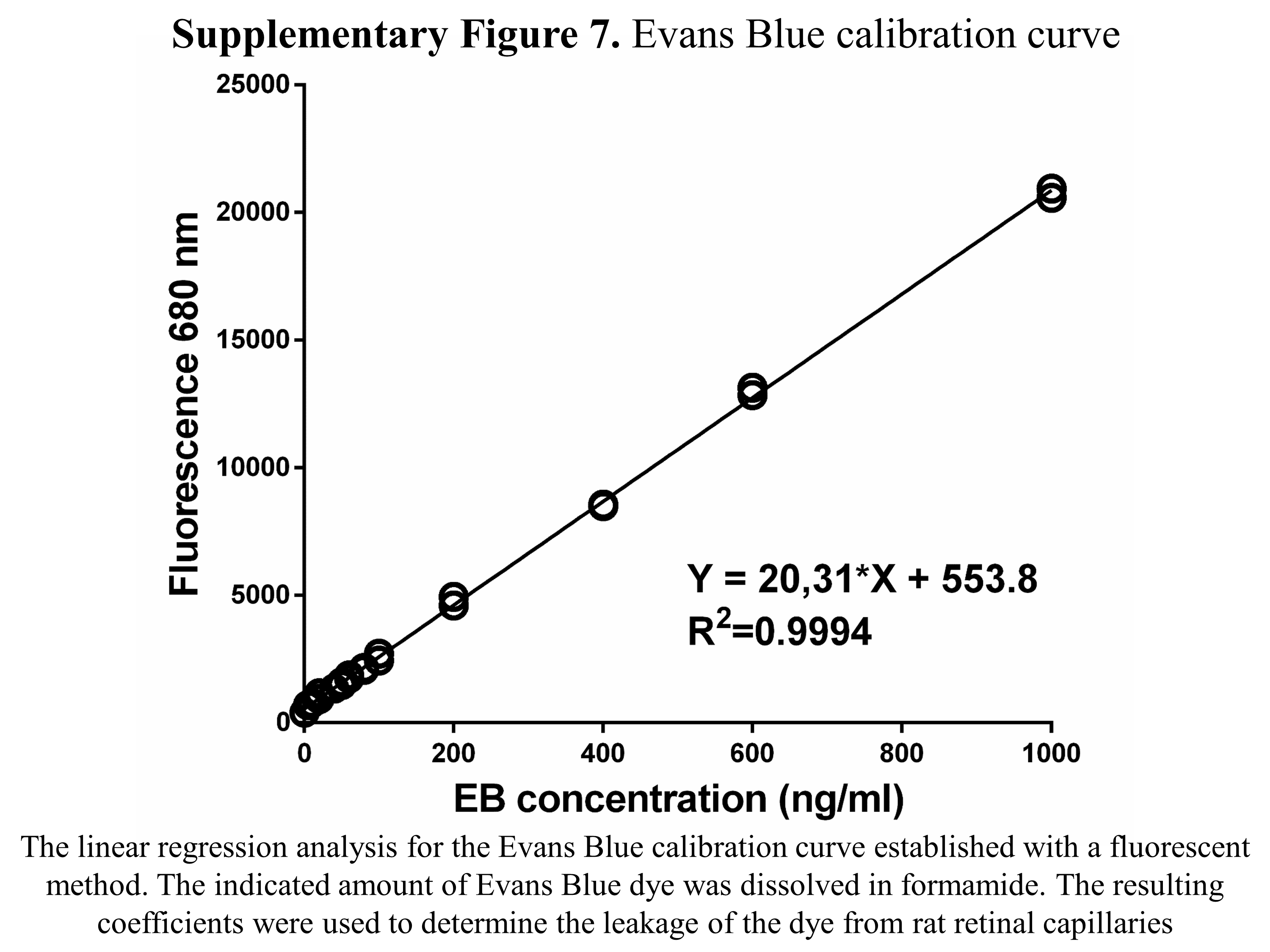

### Effect of ABCG2 inhibition on the efficiency of EYE1118

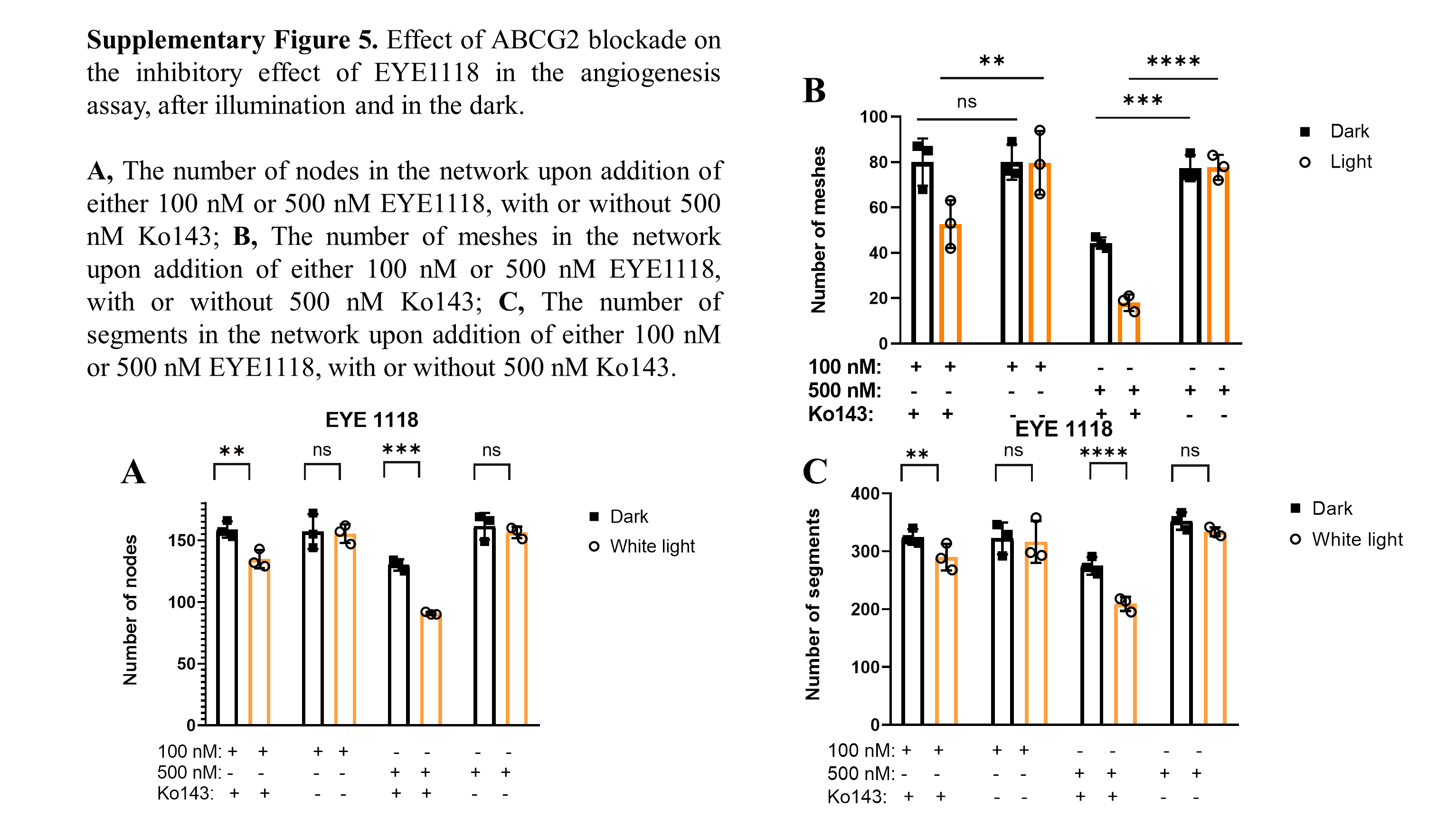

### Effect of EYE1118 on VEGFR2 phosphorylation at Y1214

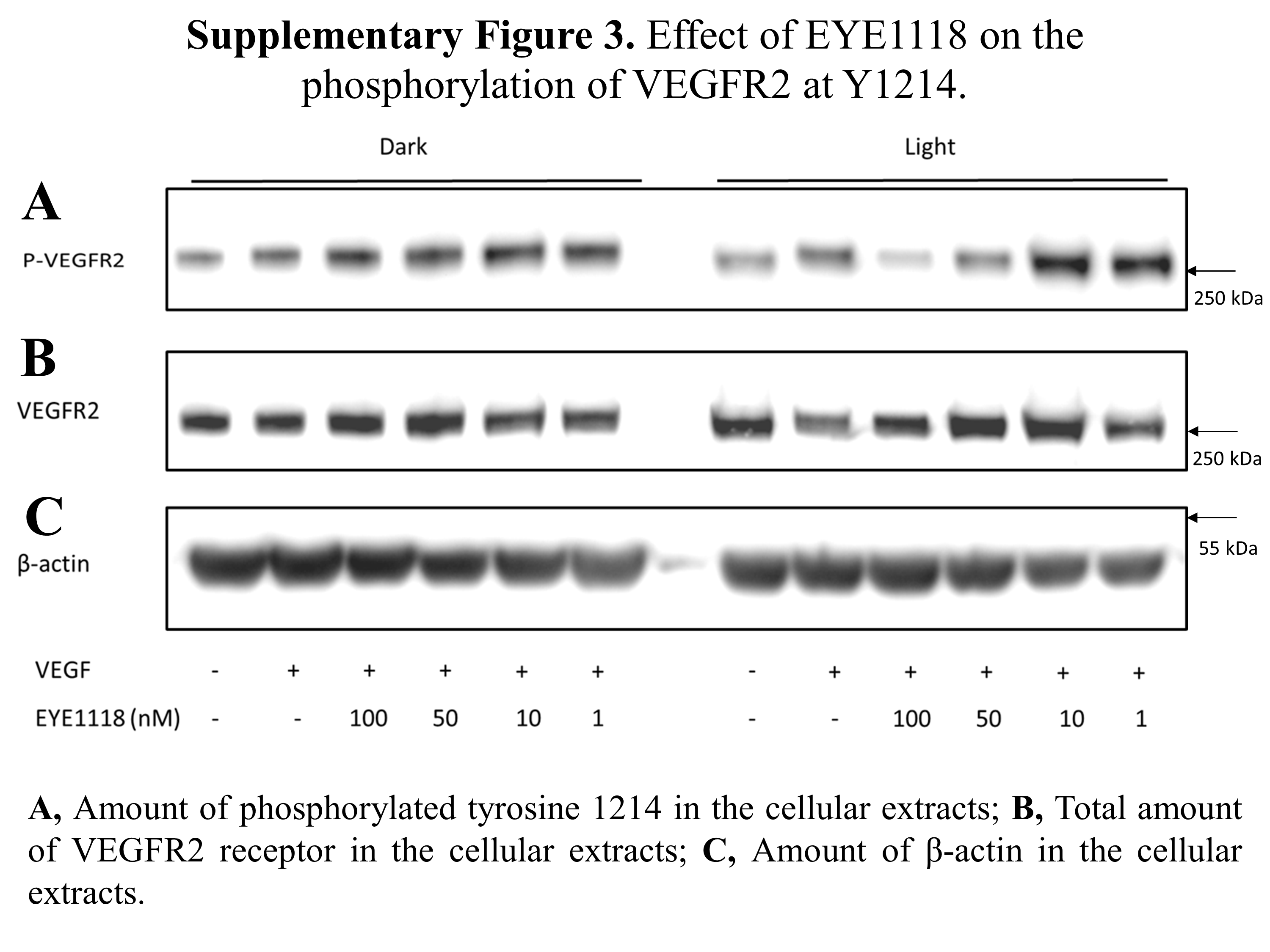

### Kinetics of VEGFR2 phosphorylation upon addition of VEGF

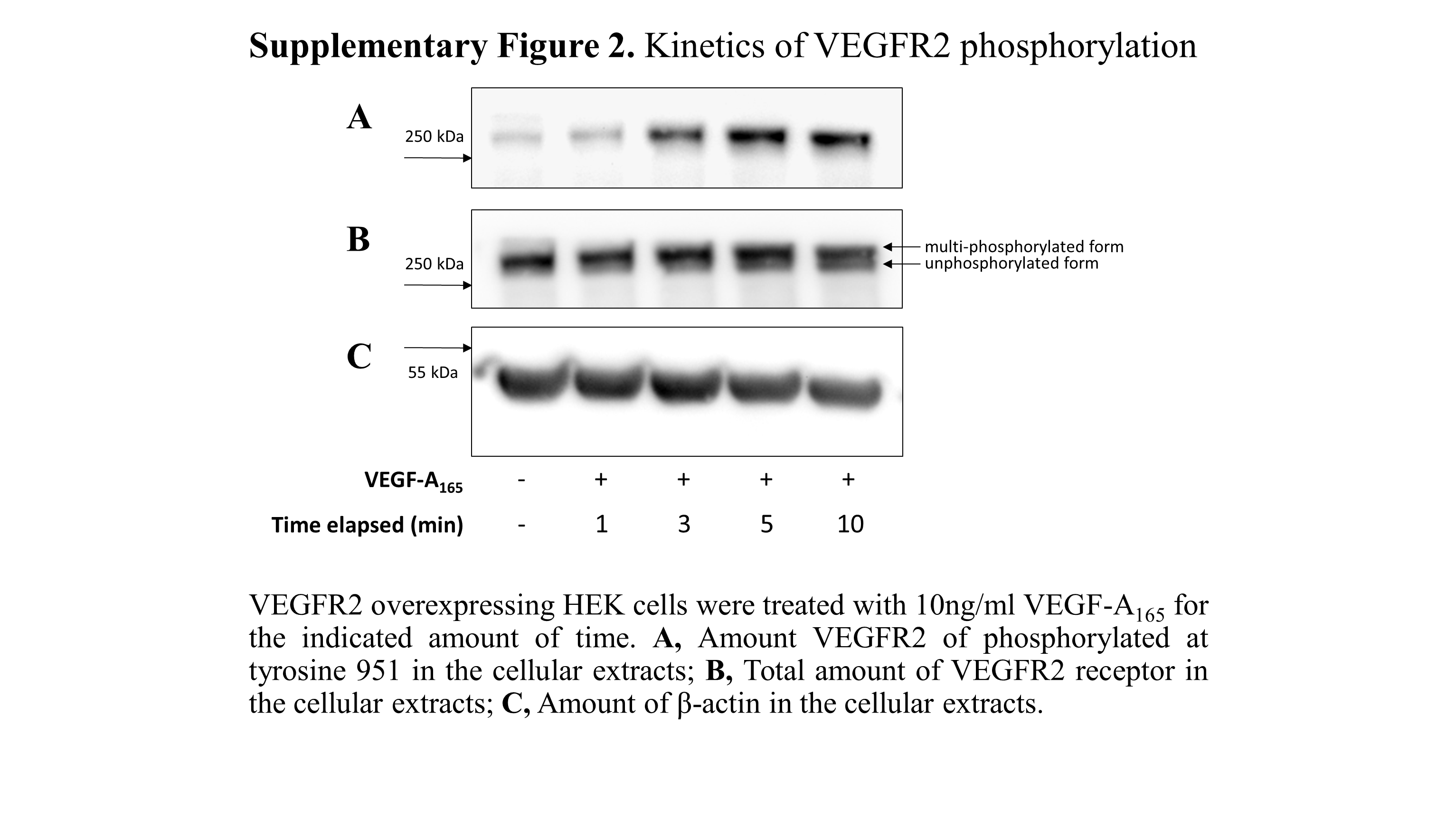

### Methodology for quantifying in vitro angiogenesis from HRMEC

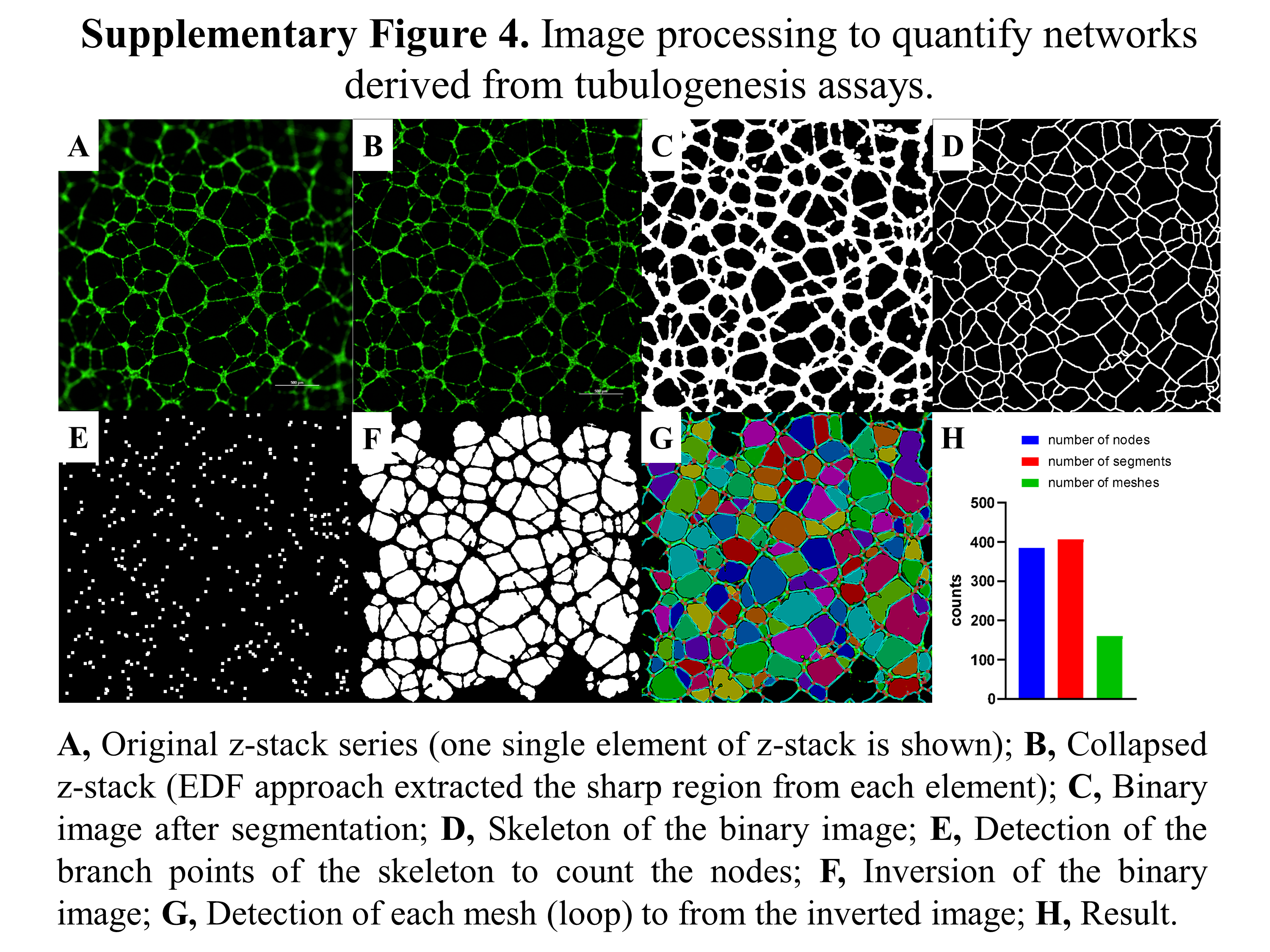

### Methodology for quantifying migration of HRMEC

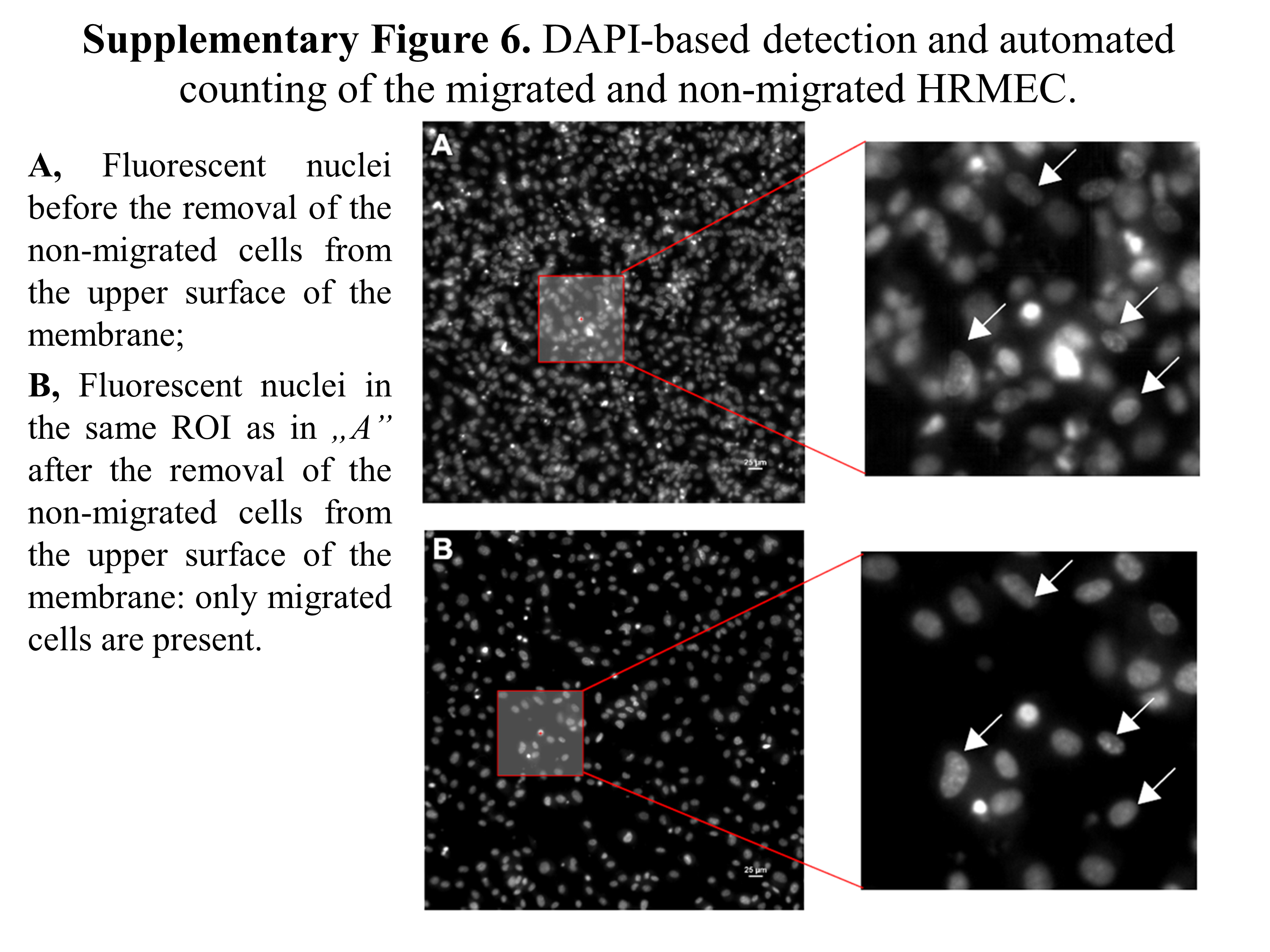

### Spectra of the light sources use in the study

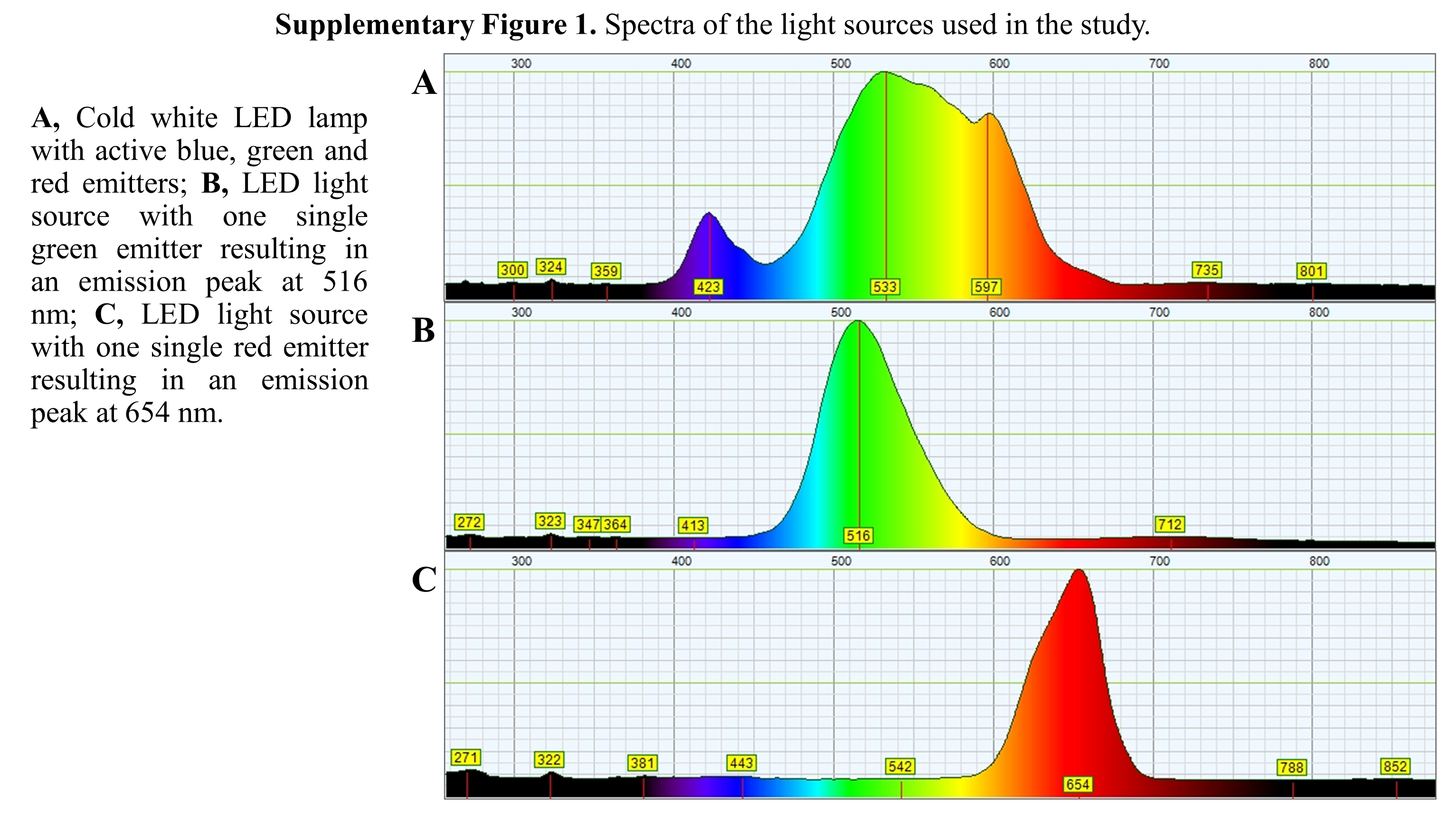
